# Decoding the physicochemical basis of taxonomy preferences in protein design models

**DOI:** 10.1101/2025.10.21.683350

**Authors:** Laura Dillon, Aaron Maiwald, Oliver Crook

## Abstract

Recent improvements in machine-learning protein design have enabled models to generate more promising candidate sequences than can be experimentally tested, placing increasing reliance on scores whose determinants remain poorly understood. Here, we profile 14 models grouped by scoring context to disentangle taxonomic and biophysical contributions. Controlling for protein family and measured biophysical properties removes 91 % to 93 % of apparent species-level variation in backbone-conditioned scores, compared with only 15 % to 29 % in sequence-only likelihoods. Species rankings also diverge: backbone-conditioned models favour archaeal and thermophilic proteins, whereas sequence-only models favour eukaryotic proteins. Backbone-conditioned scores instead reflect compactness, packing and charge, with these biases propagating into redesigned sequences. Further training ProteinMPNN on extremophile secretomes alters designed surface chemistry with minimal structural cost but largely preserves species rankings. Together, these findings show how scoring context and training data shape what protein design models favour, with implications for which sequences are generated and prioritised for experimental testing.

## Introduction

The central challenge in protein engineering is determining which amino acid sequences will reliably fold into stable structures and carry out specific functions. This sequence to structure to function relationship underlies enzyme catalysis, signalling, and immunity, but the astronomical size of sequence space makes direct exploration infeasible ^1–3^. For much of the last century, the protein folding problem ^4,5^ remained largely intractable ^6^. To design new proteins, researchers modified existing structures through rational design and directed evolution, or relied on computationally expensive physics-based energy functions ^7–12^.

Machine-learning models have shifted protein engineering from local, hypothesis-driven modification toward global, data-driven exploration^7,13–20^. Sequence-only language models estimate sequence plausibility from large protein databases, and backbone-conditioned models estimate which sequences are compatible with a given backbone. These models have designed sequences that are experimentally validated to fold and function as intended, but their scores outputted during the design process are sometimes misleading proxies of protein fitness. A high-scoring sequence may reflect functional constraints, structural compatibilities, evolutionary frequencies, taxonomic representations, or a training-derived shortcuts, but these different sources of preference are collapsed into a single score ^21^.

The gap between model metrics and protein fitness creates an evaluation challenge for protein design. Standard metrics such as perplexity, likelihood, and sequence recovery summarise fit to training-like data, but they do not reveal which biological or dataset-derived features drive model preference ^22^. Analogous issues occur across machine learning, from computer vision ^23^ to natural language processing ^24–27^, such as demographic disparities in facial recognition^28^ and inequities reinforced by healthcare algorithms^29^.

For protein design, we define bias as a systematic, reproducible preference in model scoring or generation that cannot be interpreted from aggregate accuracy alone. Bias may reflect training-data composition, architectural constraints, or learned biophysical regularities rather than target-specific functional fitness. Unlike vision or language, where users can quickly spot problematic outputs, protein design biases typically remain hidden until resource-intensive experimental validation reveals them. However, bias acts both as a possible failure mode and a diagnostic signal: a way to ask what a model has learned to favour, whether that preference is biologically interpretable, and whether it can be redirected.

Previous work has shown that protein language models carry taxonomic preferences. Ding & Steinhardt (2024) found that protein language models assign higher likelihoods to proteins from model organisms such as *E. coli* and *H. sapiens*, penalising functionally comparable proteins from under-represented species ^30^. Shaw et al. (2023) similarly found that protein language models converge on high-frequency amino-acid patterns, under-sampling rare residues and motifs^31^. These findings establish that sequence databases can imprint systematic preferences on learned sequence models, but they leave open whether the same mechanism applies to backbone-conditioned design models.

Backbone-conditioned models could behave differently than protein language models as they condition on three-dimensional backbone geometry rather than sequence alone (Fig. 1). Since structure is often more conserved than sequence ^32–34^, models such as ProteinMPNN ^35^, ESM-IF^36^, and MIF^37^ might learn more generalisable constraints on fold-compatible sequences. Yet structural databases carry their own skews: the Protein Data Bank (PDB) over-represents proteins that are experimentally tractable, rigid, stable, and crystallisable ^38,39^, while under-representing membrane proteins^40,41^ and intrinsically disordered regions^42–44^. Backbone-conditioned models may therefore replace one form of preference with another, shifting from taxonomic sequence statistics toward biophysical features enriched in structural databases.

**Figure 1.**
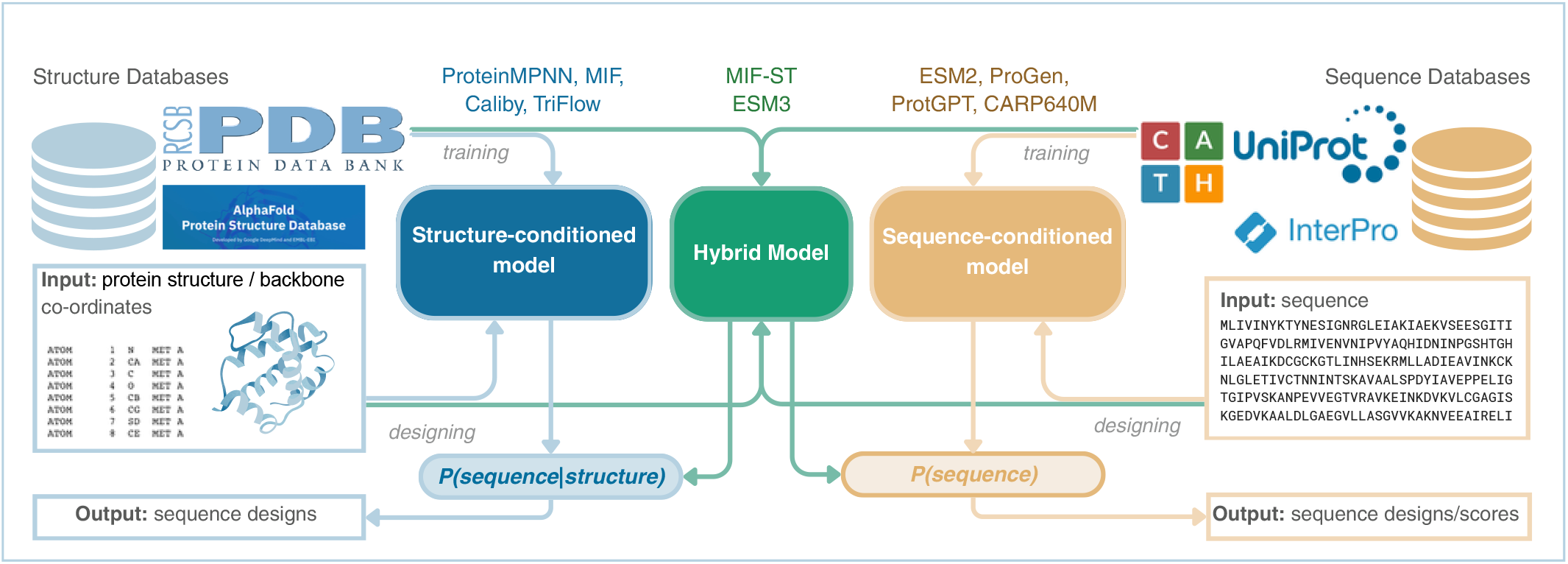
Data modalities and conditioning in protein models. Structure-conditioned models (blue box; e.g. ProteinMPNN and ESM-IF) use protein backbone information to model *P* (sequence *|* structure). Sequence-only models (orange box; e.g. ESM2 and CARP-640M) model *P* (sequence) from sequence databases. Hybrid models (green box) combine structural and sequence-derived information: MIF-ST incorporates pretrained sequence representations into structure-conditioned inference, whereas ESM3 supports conditioning on sequence and structure tracks. Training-corpus composition and information supplied during scoring are distinct dimensions that can contribute to taxonomic and biophysical preferences.

Here, we compare 14 protein design models that differ in data modality, training corpus, architecture, and scoring context. For quantitative summaries, we group them by the information available during scoring: backbone-conditioned, structure plus native-sequence context, or sequence-only. We combine variance decomposition, species-level preference rankings, biophysical preference landscapes, redesign experiments, and fine-tuning on ecologically selected extremophile secretomes to disentangle the sources of model preference and link them to generated sequence properties.

## Results

### Backbone-conditioned scores retain little taxonomic variance

We first asked whether apparent taxonomic preferences persist after accounting for protein family and measurable biophysical properties. We analysed a dataset of 10,146 proteins from 495 species and 281 families, augmenting a dataset of Ding & Steinhardt (2024) with additional archaeal extremophiles and under-represented families^30^. The protein dataset covers the three domains of life: eukaryotes (organisms with membrane-bound nuclei, including humans, yeast, and plants), bacteria (single-celled organisms without nuclei, such as *E. coli* ), and archaea (single-celled organisms often found in extreme environments such as hot springs). This dataset provides broad evolutionary coverage, sampling proteins from thermophilic organisms thriving at high temperatures, mesophilic organisms adapted to moderate conditions, and species ranging from microscopic prokaryotes to complex multicellular organisms.

We evaluated the ProGen2 model family and report XL as its representative in the primary comparisons. The base checkpoint is retained separately in supplementary sensitivity outputs. We compared fourteen representative models that differ in data modality, training corpus, architecture, and scoring context (Supplementary Table S2). For quantitative comparisons, we grouped them according to the information available when scoring each residue (Supplementary Table S2; Table 1). Backbone-conditioned models assess sequence compatibility with a supplied structure or backbone without access to the complete surrounding native sequence (e.g. ProteinMPNN on Protein Data Bank structures). Models scored with structure plus native-sequence context use both structural information and visible surrounding residues when scoring a masked position (ESM3, and MIF on CATH: a hierarchically organised database of approximately 19,000 protein domain structures classified by fold similarity). Sequence-only models score proteins without supplied structure (ESM2 on UniRef90/50, CARP-640M on UniRef50, ProGen2-XL, and ProtGPT2).^19,45–51^.

**Table 1.** Apparent species signal is largely removed by family and biophysical adjustment in backbone-conditioned models, but retained in several sequence-only language models. Species-only *R*^2^ measures the apparent taxonomic signal before adjustment. Residual species Δ_spec_ measures the remaining species contribution after controlling for protein family and 14 measured biophysical features. Attenuation is the percentage of the species-only signal removed by this adjustment. Retention measures how much per-species effect remains within protein families. Full model-level values are reported in Supplementary Table S6.

| Scoring context | Models included | Species-only $R^2$ (%) | Residual $\Delta_{\text{spec}}$ (%) | Atten. (%) | Within-family retention |
| --- | --- | --- | --- | --- | --- |
| Backbone-conditioned | ProteinMPNN; SolubleMPNN; Caliby; SolubleCaliby; ESM-IF; TriFlow | 20.9–33.9 | 1.6–2.8 | 91–93 | 0.22–0.28 |
| Structure + native sequence context | MIF; MIF-ST; ESM3-structure | 23.5–34.4 | 3.4–15.9 | 33–87 | 0.31–0.75 |
| Sequence-only | ESM3-sequence; ESM2-15B; CARP-640M; ProGen2-XL; ProtGPT2 | 18.0–46.6 | 5.2–34.5 | 15–84 | 0.38–0.82 |

The model outputs are per-residue compatibility scores, oriented so that higher values indicate stronger model preference for the wild-type sequence in its scoring context. We decomposed each model’s score variance using nested ordinary-least-squares models containing species, protein family, and 14 measured biophysical features (Methods). Protein family is a necessary control because families differ in function, composition, and taxonomic distribution, making raw species effects difficult to interpret.

After controlling for family and biophysical features, residual species variance was consistently small for backbone-conditioned models but remained larger in sequence-only language models (Table 1). Species alone explained a comparable fraction of score variance across scoring contexts before adjustment, indicating that raw species effects are not by themselves diagnostic of residual taxonomic bias. The distinction appeared after adjustment: 91 % to 93 % of the apparent species signal was removed in backbone-conditioned models, whereas only 15 % to 29 % was removed in the sequence-only language models ESM2-15B, CARP-640M, ProGen2-XL, and ProtGPT2. A within-family check supported the same interpretation, with low species-effect retention in backbone-conditioned models and high retention in classical sequence-only language models.

Models with visible native-sequence context occupied intermediate or boundary positions. MIF and ESM3-structure retained relatively little residual species signal after adjustment, whereas MIF-ST retained substantially more, consistent with its additional sequence-derived representations^37^. ESM3-sequence was also a boundary case within the sequence-only scoring context: although it was scored without supplied structure, its residual species signal was much smaller than that of the sequence-only language models. Thus, the apparent taxonomic signal in backbone-conditioned scores is largely explained by family composition and measured biophysical properties, whereas several sequence-only likelihoods retain stronger organism-level dependence within families. The separation was robust to removing ribosomal proteins and to domain-balancing controls (Supplementary Section S3.4).

The unique contribution of measured biophysical features to score variance is summarised in Supplementary Table S24. These results clarify how model preferences differ across scoring contexts. Backbone-conditioned scores are more strongly organised by measurable biophysical compatibility, while classical sequence-only likelihoods retain a larger residual component associated with organism identity after family and measured biophysics are accounted for. We next ask whether these components have consistent direction.

### Taxonomic preference differs across scoring contexts

We next asked whether the remaining taxonomic signal favours the same regions of the tree of life across model classes. To quantify directional taxonomic preferences (Fig. 2), we used a species-level Elo rating adapted from chess rankings^52^, following Ding & Steinhardt (2024) ^30^. A pLDDT-residualised variant is reported as a robustness check (Supplementary Table S10).

**Figure 2.**
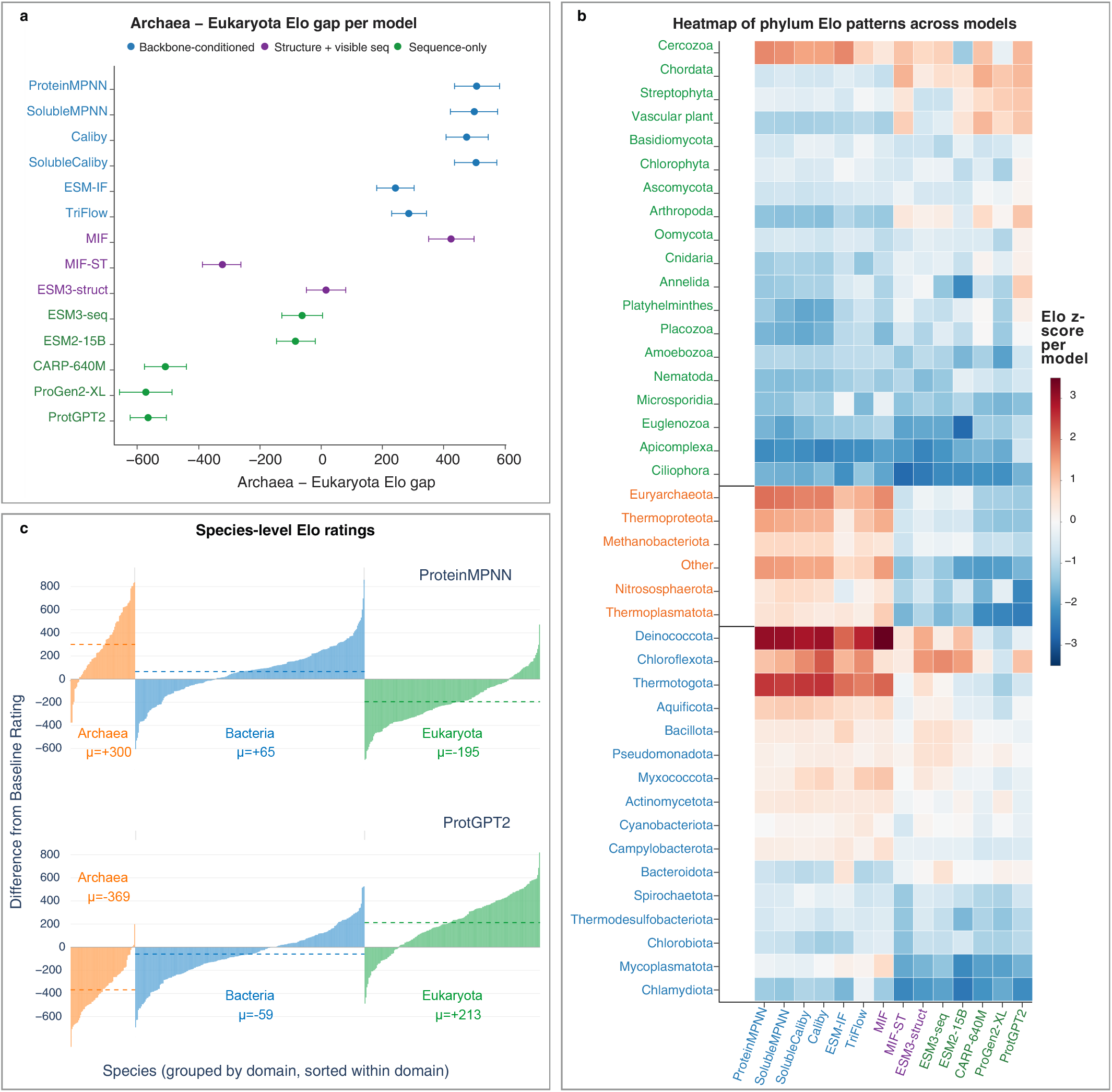
Taxonomic preference direction inverts between training modalities. Species-level Elo ratings (*∼* 1500 baseline; 495 species overall (492 for ESM2), 281 protein families, 50 permutations, unweighted) were calculated from within-family pairwise comparisons across all 14 models. A rating above 1500 means that a model tends to score that species’ proteins above the family average. **a,** Archaea–Eukaryota Elo gap for each model, calculated as the mean Archaea rating minus the mean Eukaryota rating, with 95% percentile intervals from 2,000 within-domain species bootstrap resamples (seed 0). Points are coloured by scoring context: backbone-conditioned (blue), structure plus visible native-sequence context (purple), and sequence-only (green). Backbone-conditioned models, MIF and ESM3-struct produce positive, Archaea-leaning gaps, whereas sequence-only models and MIF-ST produce negative, Eukaryota-leaning gaps. ESM3-struct lies near zero. **b,** Heatmap of species-level Elo z-scores averaged within each phylum. Ratings are first standardised across species within each model using the population standard deviation, then averaged with equal species weights. Phyla are grouped by domain, with row-label colours indicating Eukaryota (green), Archaea (orange), and Bacteria (blue). Models are ordered by scoring context. Warm cells indicate phyla that a model rates above its own mean. Most backbone-conditioned models concentrate preference on thermophilic prokaryotes, whereas sequence-only models shift towards eukaryotic phyla. Saturation represents each model’s standardised species preferences rather than its overall taxonomic effect. Other denotes unassigned archaeal lineages. **c,** Per-species Elo ratings relative to baseline for ProteinMPNN and ProtGPT2. Species are grouped by domain and sorted within each domain; dashed lines mark the domain means. ProteinMPNN rates Archaea highest and Eukaryota lowest, whereas ProtGPT2 shows the opposite pattern, illustrating the inversion in panel a. Source data are provided as a Source Data file.

Elo ratings update based on head-to-head performance comparisons according to:

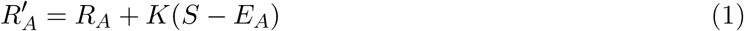

where *R_A_* is species A’s current rating, *K* = 32 is the update magnitude, *S* is the actual outcome (1 if species A wins, 0 if it loses), and *E_A_* = 1*/*(1 + 10^(^*^RB −RA^*^)^*^/^*^400^) is the expected outcome based on current ratings. If a model scores a protein from species A higher than one from species B within the same protein family, species A gains rating points while species B loses points, with larger changes occurring when outcomes go against expectations (e.g., when a low-rated species defeats a high-rated species).

Within each protein family, we conducted pairwise comparisons ranking species by model scoring preferences. Most backbone-conditioned models assigned higher scores to archaeal species, producing positive Archaea–Eukaryota gaps on a *∼*1500 Elo scale (ProteinMPNN +496, SolubleMPNN +488, Caliby +463, SolubleCaliby +494, TriFlow +273; Table 2, with full per-domain statistics in Supplementary Tables S11 and S12). In contrast, pure sequence language models favoured eukaryotic species most strongly (ProtGPT2 *−*581, CARP-640M *−*525, ProGen2-XL *−*589). Models scored with structure plus native-sequence context were heterogeneous: MIF favoured Archaea, MIF-ST favoured Eukaryota, and ESM3-structure lay close to zero on the Archaea–Eukaryota contrast.

**Table 2.** Mean species Elo by domain (*∼*1500 baseline; 495 species, 281 families, 50 permutations) and the Archaea*−*Eukaryota gap. Ratings and domain means are unweighted; ESM2 has finite ratings for 492 species. Values are derived model estimates rather than independent replicate measurements. Source data are provided as a Source Data file.

| Model | Scoring context | Archaea | Bacteria | Eukaryota | Arch–Euk | top |
| --- | --- | --- | --- | --- | --- | --- |
| ProteinMPNN | Backbone | 1800 | 1565 | 1305 | +496 | Archaea |
| SolubleMPNN | Backbone | 1800 | 1559 | 1312 | +488 | Archaea |
| Caliby | Backbone | 1770 | 1572 | 1307 | +463 | Archaea |
| SolubleCaliby | Backbone | 1783 | 1582 | 1289 | +494 | Archaea |
| ESM-IF | Backbone | 1581 | 1590 | 1352 | +229 | Bacteria |
| TriFlow | Backbone | 1660 | 1541 | 1387 | +273 | Archaea |
| MIF | Struct.+seq. | 1731 | 1573 | 1319 | +412 | Archaea |
| MIF-ST | Struct.+seq. | 1278 | 1474 | 1615 | –338 | Eukaryota |
| ESM3-structure | Struct.+seq. | 1415 | 1590 | 1413 | +2 | Bacteria |
| ESM3-sequence | Sequence-only | 1358 | 1591 | 1434 | –76 | Bacteria |
| ESM2-15B | Sequence-only | 1399 | 1531 | 1497 | –98 | Bacteria |
| CARP-640M | Sequence-only | 1154 | 1460 | 1679 | –525 | Eukaryota |
| ProGen2-XL | Sequence-only | 1066 | 1504 | 1655 | –589 | Eukaryota |
| ProtGPT2 | Sequence-only | 1131 | 1441 | 1713 | –581 | Eukaryota |

The contrast between MIF and MIF-ST provides a useful internal comparison. Both models use a masked inverse-folding architecture, but MIF-ST additionally incorporates sequence pre-training through CARP-640M residue-level embeddings. MIF showed the archaeal-leaning pattern observed in most backbone-conditioned models, favouring Archaea (+412), whereas MIF-ST showed the sequence-like pattern, favouring Eukaryota (*−*338). The inversion between MIF and MIF-ST indicates that sequence pre-training can substantially alter taxonomic preference direction, even when structural information remains available.

Beyond domain-level trends, the top-ranked species for each model are interpretable given its training data (Supplementary Table S13). Models with structural input (ProteinMPNN, ESM-IF, Caliby, TriFlow, and MIF) rank proteins from thermophilic prokaryotes highest, including archaeal hyperthermophiles and thermophilic bacteria (*Pyrococcus*, *Thermococcus* and *Thermus thermophilus*), consistent with a preference for the compact, well-packed folds that dominate experimental structure databases. Closely matched models diverge in which taxa supply these folds: ProteinMPNN’s top ten is 8/10 archaeal whereas ESM-IF’s is 8/10 bacterial. Despite highly correlated rankings (Spearman *ρ* = 0.72) they share only 20% of their top-ten species. In contrast, models trained on UniRef sequences inherit that corpus’s taxonomic skew: the sequence language models (ESM2-15B and ESM3) favour model and disease-relevant bacteria (*E. coli*, *Salmonella*, *Yersinia*, *Klebsiella*), while ProGen2-XL ranks predominantly plant and animal species (nine of its top ten are eukaryotic). CARP-640M and ProtGPT2 shift towards mammalian and domesticated species (*Homo sapiens*, *B. taurus*, *O. aries*, *C. lupus*), a pattern also seen in MIF-ST, plausibly reflecting its CARP-derived sequence pre-training on UniRef50. In every case the favoured taxa mirror what is abundant in the model’s training data ^53,54^.

### Backbone-conditioned scores track biophysical properties

To identify what replaces taxonomic dependence in backbone-conditioned models, we mapped each model’s scores onto a low-dimensional biophysical feature space. The first two principal components of the 14 biophysical features account for 40.0 % of feature variance (feature loadings, Supplementary Table S14). The three domains represented in the protein dataset do not form separate clusters. Quantifying the per-domain distributions by the Bhattacharyya overlap of their bivariate Gaussians, all pairs overlap heavily (Bacteria–Archaea 95 %, Eukaryota–Archaea 91 %, Eukaryota–Bacteria 84 %), and the PC1 means are Bacteria *−*0.66, Archaea *−*0.17, Eukaryota +0.66. PC1 broadly corresponds to surface charge, chain length and hydrophobicity. Negative PC1 proteins have high isoelectric point, high basic-residue fraction, thus are mostly small, basic, positively-charged proteins (ribosomal-like, nucleic-acid-binding character). Positive PC1 proteins have longer chains, larger C*β* spread, and higher GRAVY, meaning they are calculated to be more acidic, longer, and more hydrophobic/aromatic. PC2 is mostly orthogonal to charge and contrasts packing and composition-related features. Negative PC2 proteins are compact, hydrophobic, ordered, well-packed folds, with smaller C*β* distances and more long-range contacts. Positive PC2 proteins have larger C*β* distances, looser packing, bulkier residues and higher sequence-based instability index. These descriptors do not measure thermal stability. As these two PCs account for only 40.0% of total variance, position in the plane is a coarse biophysical fingerprint rather than a complete description. Figure 3b shows the protein dataset projected into the PCA space, showing where well characterised proteins are placed. Bacteria occupy the most negative-PC1 (compact and positive charge) region; eukaryotic proteins extend somewhat further toward large, hydrophobic values. The complete set of per-model GAM preference landscapes is shown in Supplementary Fig. S7.

**Figure 3.**
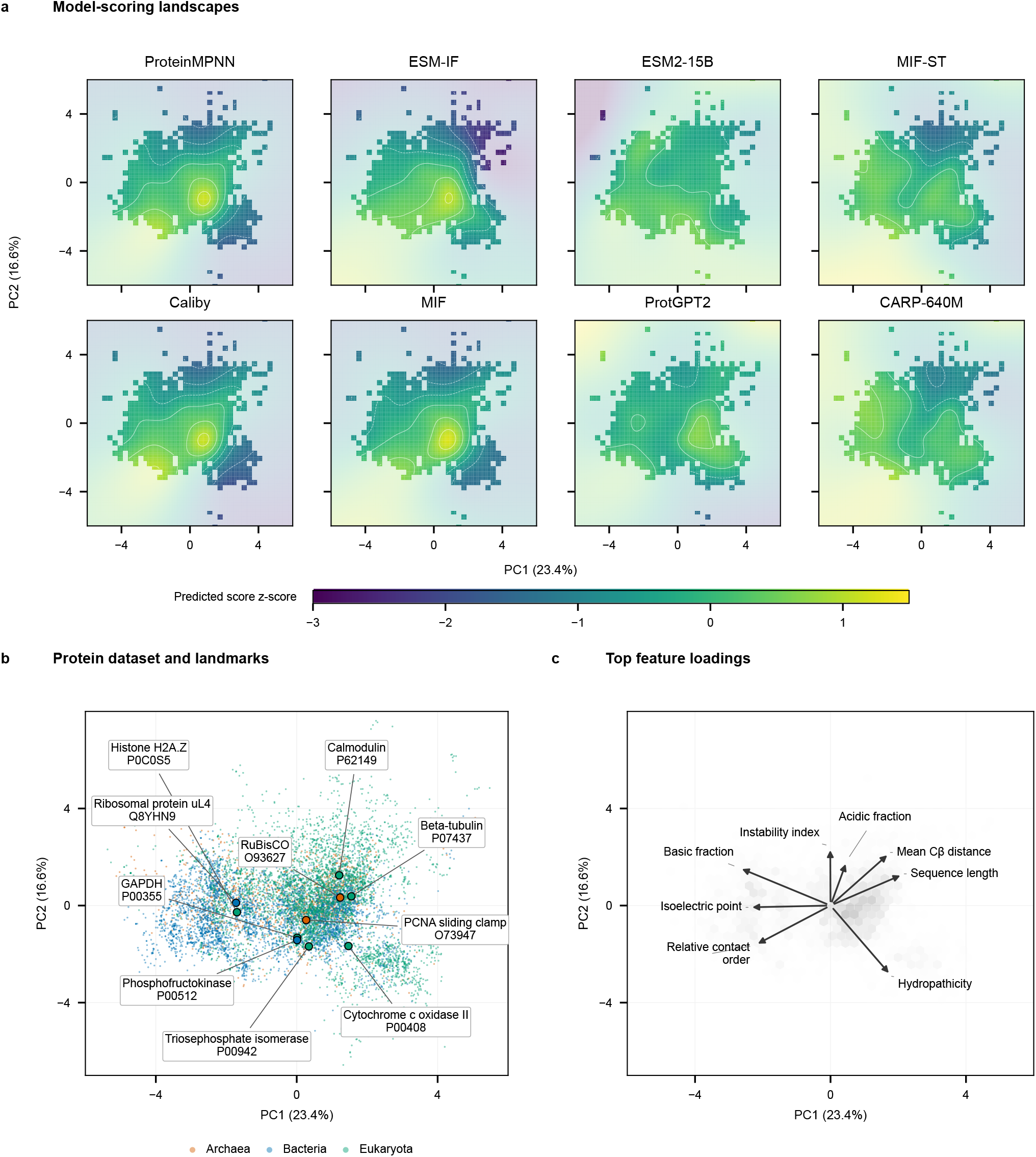
Biophysical organisation of the protein dataset and model-scoring landscapes. a, Generalised additive model (GAM) landscapes show within-model z-scored scores in the 14-feature principal-component analysis (PCA) space. Purple to yellow indicates increasing predicted score; contours connect equal values. Cells with fewer than three supporting proteins or outside the observed convex hull are shown at 22% opacity as background extrapolation; full colour and contours identify supported predictions. All panels share a colour scale from *−*3 to +1.5. **b,** Protein dataset projected onto the same two components, coloured by taxonomic domain: Archaea (orange), Bacteria (blue), and Eukaryota (green). Selected proteins are labelled. **c,** Selected feature loadings. Arrow direction indicates each feature’s association with the components; length indicates its contribution within the displayed plane, with a common display scaling factor for all eight arrows. Negative PC1 is associated with higher isoelectric point, basic-residue fraction and relative contact order; positive PC1 with longer, more acidic and more hydrophobic proteins. PC2 contrasts instability and mean C*β* distance with hydrophobicity and contact order. Source data include all coordinates, the ten landmark identifiers and the loading-arrow scale. Source data are provided as a Source Data file.

We fitted a generalised additive model (GAM) of each model’s score over that plane (Methods). The GAM deviance explained asks how much a given model’s scoring behaviour is captured in the features, which ranges 9 % to 52 % across models and is largest for backbone-conditioned models (ESM-IF *≈* 52 %, SolubleMPNN *≈* 50 %, ProteinMPNN *≈* 44 %) and smallest for the sequence-only ESM2-15B (*≈* 9 %); the representative ProGen2-XL scores give *≈* 9 % (Supplementary Table S15). The split between backbone-conditioned models and sequence models tracks with the relative influence biophysical features appear to have on each class. Backbone-conditioned models place their highest scores in the compact, well-packed region of the plane; sequence-only models show weaker preferences. However, the domains share a region of biophysical space with substantial overlap, and the backbone-conditioned models’ compactness preference maps onto this shared region rather than onto cleanly separated taxonomic clusters.

### Designed sequences show model-specific biophysical shifts

To see whether scoring preferences also shape generated sequences (Fig. 4c), we redesigned 25 proteins with seven models (ProteinMPNN, SolubleMPNN, Caliby, SolubleCaliby, ESM-IF, MIF, and MIF-ST), generating eight designs per protein template at sampling temperature 0.1. The template proteins selected spanned the three domains and represent a range of size and function (Supplementary Table S19). For each model and template, we averaged the eight designs, subtracted the wild-type property value, and tested paired shifts across the 14 biophysical properties of the PCA feature set using *d_z_* = mean(Δ)*/* sd(Δ), Wilcoxon signed-rank tests, and Benjamini–Hochberg FDR correction (Methods). Two of these properties (chain length and surface exposure) are invariant under fixed-backbone redesign and carry no shift, leaving 12 properties with measurable changes (Supplementary Table S20).

**Figure 4.**
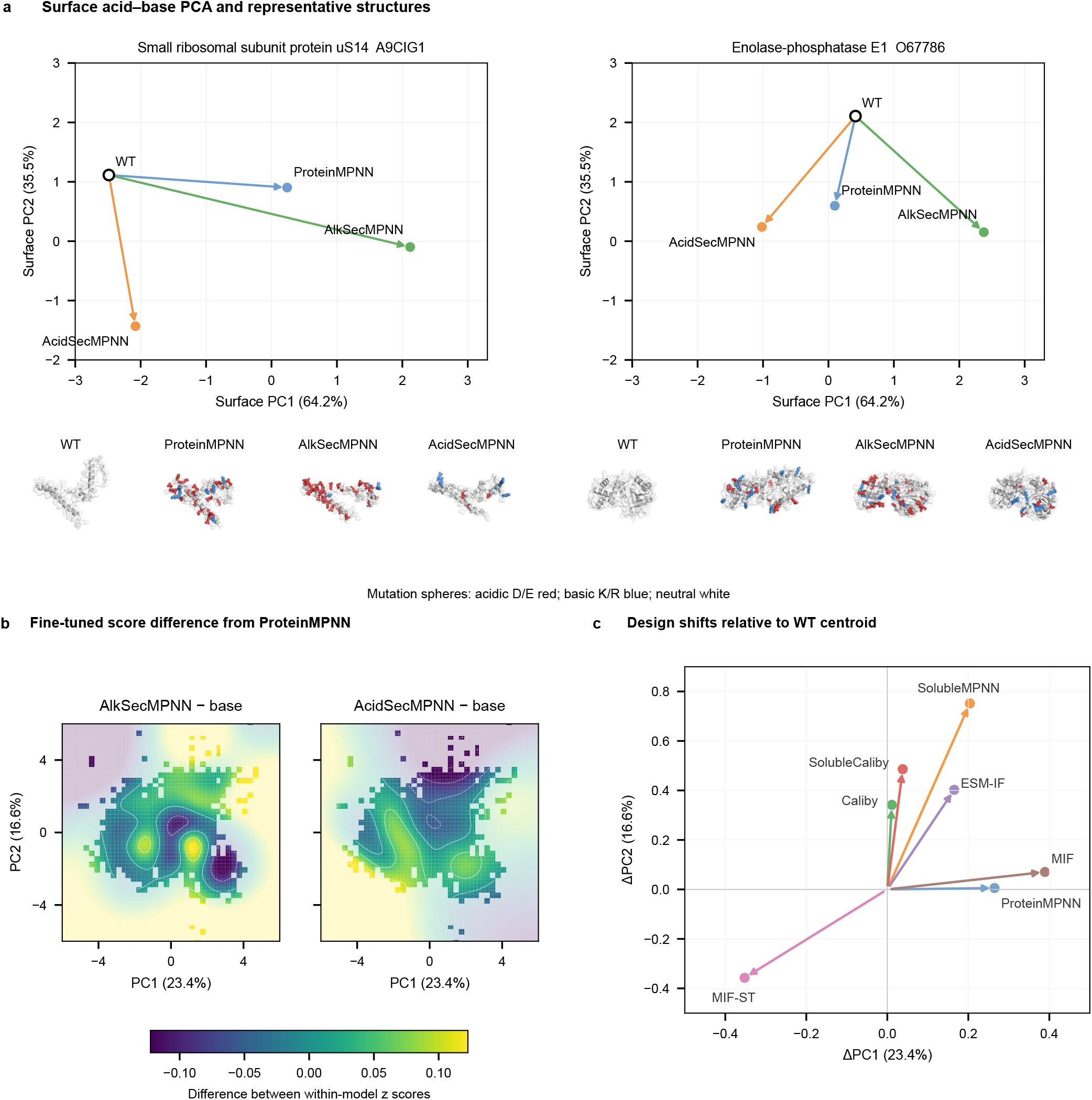
Design and fine-tuning alter biophysical preferences. a, Wild-type (WT)-to-design movement for two illustrative templates in the four-feature surface acid–base PCA used for Supplementary Table S26 (PC1, 64.2%; PC2, 35.5%). Open circles show WT; arrows end at the mean of eight designs for base ProteinMPNN (blue), AlkSecMPNN (green), and AcidSecMPNN (orange). The fit includes 1,225 WT/design records across primary and sensitivity checkpoints (Methods). Structure renders show WT and the highest-self-consistency-TM-score design among the eight samples for each model. Mutation spheres indicate the resulting residue class: acidic D/E (red), basic K/R (blue), or neutral (white). All-template surface shifts are shown in Supplementary Fig. S10. **b,** Generalised additive model landscapes of fine-tuned-minus-base scores in the separate, fixed 14-feature global natural-protein PCA (PC1, 23.4%; PC2, 16.6%). Colours show the difference between each fine-tuned model score and the base ProteinMPNN score after within-model z-scoring, from lower differences in purple to higher differences in yellow. Positive values indicate regions scored more favourably by the fine-tuned model relative to base; negative values indicate regions scored less favourably. Both panels share a colour scale capped symmetrically at the larger of their 95th percentiles of absolute fitted values in supported cells. Cells with fewer than three supporting proteins or outside the observed convex hull are shown at 22% opacity as background extrapolation; contours are restricted to supported predictions. **c,** Mean WT-to-design shifts projected into the biophysical PCA space for the seven original design models. Arrows run from the WT centroid to the centroid of the corresponding designed sequences, expressed as changes in PC1 and PC2 relative to WT. Most models shift towards positive PC2, with varying magnitude and direction. MIF-ST instead moves towards negative PC1 and PC2. Model labels identify the arrows. Source data are provided as a Source Data file.

Generated sequences exhibit model-specific biophysical tendencies that are modulated by template characteristics, demonstrating that design behaviour reflects both biases and template-specific optimisation. Five properties shift significantly (FDR*<* 0.05 in *≥* 4*/*6 structural-input models): isoelectric point, acidic-residue fraction, aromaticity, proline fraction, and helix*−*sheet contrast. The structural-input models share a broadly consistent signature: designs move toward more acidic compositions (higher acidic-residue fraction, lower isoelectric point, thus lower net charge), more proline, higher helix*−*sheet contrast, and lower aromaticity, with ProteinMPNN and Caliby additionally raising relative contact order. The structural-input models have positively correlated profiles across the 12 paired standardised property effects (*r* = 0.43–0.99), whereas MIF-ST is negatively correlated with every other model (*r* = *−*0.20 to *−*0.44), including MIF (*r* = *−*0.20). MIF-ST therefore produces a distinct biophysical response under this design protocol. The per-property design shifts are listed in Supplementary Table S20. We also tested whether the models differentially preserve annotated functional residues. Across 15 templates with curated UniProt active-site, binding, and metal-binding annotations, no model recovers functional positions significantly above background, and the only significant effect is MIF-ST recovering them below background (Δ = *−*0.158, *p* = 1.8 *×* 10*^−^*^4^; Supplementary Table S25; Supplementary Fig. S9). Differential handling of catalytic or ligand-binding residues therefore does not account for the biophysical shifts.

### Fine-tuning redirects surface acid–base properties

To test whether these preferences are malleable based on training data composition (Fig. 4a,b), we fine-tuned ProteinMPNN on selected extremophile secretomes: proteins secreted by organisms adapted to alkaline (pH *∼*9–11) or acidic (pH *∼*1–4) environments, each matched 1:1 to neutralophile-secretome controls (Methods). Secreted proteins are exposed to extracellular pH, making them a tractable setting in which organismal ecology may enrich surface chemistry without selecting proteins directly by charge. The cohorts are selected by organismal growth pH, not by the charge of individual proteins, meaning the resulting shifts in designs are likely mediated by the sequence and surface composition the model sees. The two arms produced Alkaliphile-Secretome-MPNN (AlkSecMPNN) and Acidophile-Secretome-MPNN (AcidSecMPNN). Each was evaluated alongside a matched neutralophile-trained checkpoint produced using the same continued-training protocol.

Relative to base ProteinMPNN, AlkSecMPNN designs moved toward more acidic, lower-charge surfaces along PC1 (ΔPC1 = +1.94), whereas AcidSecMPNN designs moved in the opposite direction (ΔPC1 = *−*1.00; Supplementary Table S26). The surface PCA uses four features: surface acidic, basic and ionisable residue fractions, and surface net charge. Six direct surface and sequence features are reported separately (Supplementary Table S27). AlkSecMPNN increased surface acidic-residue fraction and decreased surface net charge, charge per residue and pI. AcidSecMPNN instead decreased surface acidic-residue fraction and increased net charge, charge per residue and pI.

On held-out neutralophile backbones, the alkaliphile-trained model and its matched neutralophile control shifted surface net charge in opposite directions, whereas the corresponding acidophile-trained model and control shifted it in the same direction (Supplementary Table S27, Panel B). The extremophile-versus-control difference was therefore larger for the alkaliphile arm (*−*0.2358) than for the acidophile arm (+0.0248). We consequently interpret the alkaliphile-associated shift more strongly and treat the acidophile result as a weaker, directionally consistent effect.

The shifts preserve design quality: native-sequence recovery changed by *−*1.4 and +0.3 percentage points for the two arms, with entropy, duplicate rate, and pairwise identity comparable to base. Single-sequence AlphaFold2 self-consistency showed no detectable structural cost for AlkSecMPNN (scTM = 0.51 vs base 0.54; Δ = *−*0.02, n.s.) and a small cost for AcidSecMPNN (scTM = 0.45; Δ = *−*0.09, *p <* 0.001); both refold better than the wild-type-sequence control (scTM = 0.38; Supplementary Fig. S11). Fine-tuning did not overwrite the taxonomic structure that we observed. The Archaea*−*Eukaryota Elo gap persisted after fine-tuning (+496 base *→* +553 AlkSecMPNN, +417 AcidSecMPNN). Fine-tuning therefore redirects one local, surface-localised design tendency while leaving ProteinMPNN’s broader taxonomic preference largely intact.

## Discussion

We characterised systematic preferences in protein design models and found that they differ across model classes in ways that reflect both scoring context and training-corpus composition. Across 14 models, apparent species preferences in backbone-conditioned scores were largely attenuated after controlling for protein family and measured biophysical features, whereas sequence-only scores retained stronger within-family taxonomic dependence. Backbone-conditioned models instead organised their scores around biophysical axes such as compactness, packing, secondary-structure content, and charge. These results show that model bias can reflect family composition, organism-level sequence statistics, measurable biophysical compatibility, or training-derived enrichment of particular protein classes.

Training-corpus composition and scoring context are distinct mechanisms. Training-corpus composition determines which protein distributions are available for a model to learn, whereas scoring context determines which structural and sequence information is available when a score is assigned. Architecture and pre-training history can further modify how these sources of information are combined.

This distinction matters because modern machine-learning design models optimise learned probability distributions rather than biological fitness directly. Classical protein design methods, such as Rosetta, were framed around physical energy functions and the search for low-energy sequence–structure pairs^12,55^. Recent machine-learning methods instead learn high-probability sequences, structures, or sequence– structure pairs from large empirical datasets^18,35,36^. These objectives are related but not equivalent: a high-probability sequence under a training distribution may reflect structural compatibility, evolutionary frequency, experimental tractability, taxonomic sampling, or annotation practice rather than optimality for a target function. Our analyses ask which of these biological and dataset-derived factors are measurable in model scores and whether they propagate into generated sequences.

Training datasets provide a likely explanation for these differences. Models with structural input draw on resources derived from the PDB, CATH, AlphaFold Database, or related backbone collections. ProteinMPNN, for example, was trained on experimentally determined PDB structures filtered by resolution and split using CATH classifications ^35^; ESM-IF used millions of AlphaFold-predicted structures from UniRef sequences ^36^; and MIF-family models combine structural conditioning with sequence-derived information in different ways^37^. These structural resources are not neutral samples of protein space. Experimentally determined structures over-represent proteins that are expressible, soluble, ordered, stable, and amenable to crystallography or cryo-EM^38,39^, while under-representing membrane proteins^40,41^, flexible regions, low-complexity sequences, and intrinsically disordered proteins ^42–44^. AlphaFold-derived structures expand coverage substantially, but they also introduce confidence-dependent filters and may carry geometric regularities inherited from prediction models and their underlying sequence resources ^53,56^. Structural-input models may therefore reduce residual organism-level sequence bias while acquiring preferences for proteins that resemble structurally tractable database entries.

Sequence-only models are exposed to a different sampling process. Protein language models learn from large sequence corpora derived from UniProt, UniRef, BFD-like collections, metagenomic resources, or related clustered datasets^14,54,57^. These databases cover a wider range of organisms than experimentally solved structures, but they still reflect sequencing effort, annotation depth, reference-proteome selection, clustering procedures, and the over-representation of model organisms, pathogens, and medically or commercially important taxa. Because sequence-only models do not condition on a fixed backbone, their likelihoods can absorb organism-level sequence statistics more directly. The larger residual species terms we observe in sequence-only models are consistent with this mode of training: after family and measured biophysical properties are accounted for, substantial taxonomic signal remains in the score.

The contrast between MIF and MIF-ST supports the role of sequence pre-training in shifting taxonomic preference. MIF follows the archaeal-leaning pattern observed in most backbone-conditioned models, whereas MIF-ST, which incorporates CARP-derived sequence representations, shifts toward the sequence-model pattern in domain preference^37^. This inversion suggests that sequence pre-training can alter the taxonomic direction of a structure-aware model. The comparison should not be read as a complete mechanistic isolation of training data, because MIF-ST also behaves as an outlier in several design analyses, but it strengthens the broader conclusion that sequence pre-training can change which sources of preference dominate (Supplementary Section S11).

Structural-input and sequence-only models encode different mixtures of biological and database-derived signal. For backbone-conditioned models, a low score is more closely associated with measured backbone-compatible biophysical properties in this dataset, but that does not mean it measures biological fitness directly. It may also reflect preferences for compact, ordered, soluble, or structurally typical proteins. For sequence-only models, likelihood captures broad evolutionary and compositional regularities, but remains more strongly entangled with organism-level sequence statistics. Model choice should therefore depend on the design task as the same preference may be useful when it aligns with the target, and limiting when it steers away from viable but under-represented protein classes.

Current validation practices may miss these systematic preferences because protein design is an end-to-end pipeline rather than a model-only process. AI-driven design is a sequence of dataset curation, model training, candidate generation, in silico filtering, experimental validation, and deployment^58^. Experimental validation remains indispensable, but candidates are commonly preselected using structure-prediction confidence, physics-based scores, clustering, diversity filters, and manual inspection. Synthesising and testing top-scoring designs can therefore show that a pipeline recovers functional proteins, but it does not reveal whether alternative regions of sequence-property space were consistently down-weighted before experimental testing.

Proteins whose useful properties differ from those enriched in training datasets are most exposed to this. Membrane proteins, flexible proteins, intrinsically disordered regions, proteins requiring unusual charge distributions, and proteins from under-represented taxa may all be systematically disfavoured if model scores or filters reward structural typicality. Derry et al.’s observation that models trained on X-ray structures can learn to disfavour lysine illustrates the general problem: a model may learn experimental tractability as if it were biological optimality ^38^. Our results extend this concern across model classes by showing that differences in data modality, training-corpus composition, and scoring context produce distinct, measurable preference structures.

The redesign and fine-tuning experiments show that model preferences affect generation and can be partially redirected. Structural-input models shifted wild-type templates toward characteristic biophysical profiles, including lower net charge, greater acidity, altered secondary-structure composition, higher contact order (Fig. 4c); these shifts correlated with the corresponding scoring preferences, indicating that properties inferred from wild-type scoring propagate into generated sequences. Model-guided exploration therefore does not sample all plausible sequences around a backbone uniformly, but navigates through a model-specific region of sequence-property space. Continued training of ProteinMPNN on selected extremophile secretomes further showed that at least one component of this preference structure is locally malleable (Fig. 4a,b). Together, variance decomposition, Elo rankings, biophysical preference landscapes, redesign experiments, and fine-tuning separate model preference into usually collapsed components. This framework profiles models independently of aggregate benchmark performance, which may not reveal which regions of design space a model will favour ^22,58^.

These distinctions become more important as protein design increasingly uses outputs from one model as inputs to another ^46,53,59,60^. Predicted structures, generated sequences, filtered design sets, and experimentally validated successes can all feed back into future datasets. If models are trained on predicted structures or sequences selected by previous models, then model-specific preferences may become reinforced and harder to distinguish from biological regularities. Whether such feedback already affects current protein design models remains uncertain, but the rapid growth of predicted-structure resources makes the issue increasingly relevant^53,56^. Dataset documentation and preference profiling should therefore accompany model development, especially when models are applied to protein classes under-represented in their training data.

One value of bias characterisation is identifying which preferences a model carries, when they support the intended design task, when they confound interpretation, and when they can be redirected^58,61^. As AI protein design becomes increasingly practical and routine in engineering workflows^18,35,62^, the evaluation question expands from whether models can produce functional designs to which regions of sequence-property space they preferentially explore. Disentangling the sources of model preference will therefore be important for selecting models, interpreting their scores, and designing interventions that steer models toward specific applications.

## Methods

### Dataset

We built a cohort of 10 146 proteins, 495 species, and 281 protein families, augmenting Ding & Steinhardt (2024) with archaeal extremophiles and under-represented families^30^. Proteins were drawn from reviewed UniProt/Swiss-Prot entries with matched models from the AlphaFold Protein Structure Database ^53^. Species were chosen to be well represented in Swiss-Prot while covering Eukaryota, Bacteria and Archaea (including archaeal extremophiles). Proteins were retained if they used the standard amino-acid alphabet and had a sufficiently confident AlphaFold structure available; under-represented families were supplemented with additional orthologs. Proteins were grouped into families using UniProt/Pfam annotations ^63^, and a breadth filter applied iteratively until stable: families required in *≥* 5 species and species represented by *≥* 2 proteins. Two entries, P19949 and Q8TFG7, were excluded from the analysis cohort. The scoring cohort spans sequence lengths 24–2531 aa (no length filter is applied to scoring). Separately, the 25 design templates are length-filtered to 50–650 aa for tractable folding and redesign.

To ensure adequate representation across all three domains of life, we augmented the initial species pool with additional archaeal species, prioritising organisms that displayed extremophile characteristics. We retained orthologous protein groupings where functional annotations were consistent across species. For proteins from archaeal extremophiles with inconsistent functional annotations, we assigned proteins to existing functional groups based on sequence similarity while maintaining phylogenetic breadth (Supplementary Table S28). Of the 281 families, 74 span all three domains, 88 span two, and 119 are found in one. Each protein carries biophysical descriptors, available model scores, a broad functional category, and annotation flags (enzyme, transmembrane, glycosylated, PDB-associated, disordered). The variance decomposition runs on the 9963 complete-case proteins with all model scores and both PCA coordinates present.

Structures are AlphaFold DB v6 monomer models (AF-<Uniprot>-F1-model_v6)^53^. AlphaFold predictions were used so that every protein, including those without an experimental structure, carries structural information and phylogenetic breadth is retained. AlphaFold confidence (pLDDT) is carried as a covariate, both to indicate how much predicted confidence might bias the scores and to filter low-confidence designs. Experimentally-determined structures from the Protein Data Bank are available for a subset of the cohort; separately, an independent cohort of experimental X-ray structures is used to test whether the conclusions hold on experimentally determined structures and resolved chain sequences, reported in the Supplement (Supplementary Section S4, Fig. S3).

### Models and scoring

Fourteen models were evaluated. For quantitative summaries, models were grouped by scoring context: backbone-conditioned models (ProteinMPNN, SolubleMPNN, Caliby, SolubleCaliby, ESM-IF, and TriFlow); models scored with structure plus native-sequence context (MIF, MIF-ST, and ESM3-structure); and sequence-only models (ESM3-sequence, ESM2-15B, CARP-640M, ProGen2-XL, and ProtGPT2). This grouping describes the information available during scoring, distinct from data modality, architecture, pre-training history, and training corpus. Each score was reduced to a per-residue mean and oriented so that higher values indicate stronger model preference. Because the score families are different mathematical objects (including autoregressive likelihood, masked pseudo-likelihood, flow-matching marginal, and Potts energy) absolute magnitudes were not compared across models; comparisons instead used per-model variance decomposition, model-specific normalisation, and cluster-level contrasts (Supplementary Section S1.3).

### Feature set

Sixteen biophysical properties are computed per protein, all non-predicted: deterministic functions of the amino-acid sequence or the AlphaFold backbone; model likelihoods and pLDDT are excluded (pLDDT enters only as a covariate). For the PCA, variance decomposition, and property-importance analyses we use a collinearity-pruned set of 14 features (9 sequence, 5 structure; Supplementary Table S29), keeping every pairwise *|r| <* 0.8 (max in-set *|r|* = 0.74, GRAVY vs basic-residue fraction): net charge at pH 7 (*|r|* = 0.85 vs pI) and small-residue fraction (*|r|* = 0.86 vs MW per residue) are dropped, in addition to the earlier collinear exclusions (charge-per-residue vs pI; compactness *r* = 1.00 vs mean C*β* distance). The design-shift analysis instead reports per-property shifts for all 16 computed properties. Features are computed with the repository’s own extractors (sequence_features.py, structural_features.py), reused unmodified so designs are computed identically to the reference cohort.

### Elo

Taxonomic preference is quantified per model with a species-level Elo rating, following Ding & Steinhardt ^30^. Sensitivity to AlphaFold confidence is assessed using pLDDT-weighted and pLDDT-residualised variants. Within each protein family, model scores are z-scored so comparisons are between proteins of the same family; pairwise “matches” between species then update ratings. For a match in which species *A*’s protein outscores species *B*’s, the expected result is

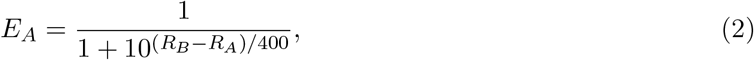

and ratings update as *R^′^_A_* = *R_A_*+*K*(*S −E_A_*) (*S* = 1 for a win, *K* = 32, baseline 1500), and symmetrically for *B*. Ratings are averaged over 50 random match orderings for stability. The full-dataset ratings cover 495 species overall: ESM2 has ratings for 492 species because the other three have no available ESM2 scores; the remaining models each have 495. For Figure 2a, 95% percentile intervals use 2,000 bootstrap resamples of species independently within Archaea and Eukaryota (seed 0), conditional on the fitted ratings; Elo matches are not refitted within these resamples.

#### Weighting variants and the pLDDT-residual control

The primary Elo is unweighted. For representative models we also calculate a pLDDT-weighted Elo, multiplying each update by the product of the two family/species mean confidence scores divided by 10,000 (bounded between 0.01 and 1), and a pLDDT-residual Elo that removes structural confidence from the score itself. For the residual control, we regress each model’s score on avg_plddt (OLS over all proteins) and recompute the unweighted Elo on the residuals (–plddt-residual). Eukaryotic AlphaFold models carry lower mean pLDDT than archaeal and bacterial ones (86.8 vs *∼*92), so a structure-quality artefact would predict the Archaea– Eukaryota gap to collapse after residualisation; the reductions differ by model, with ProteinMPNN and ESM-IF retaining positive Archaea–Eukaryota gaps and ESM2-15B retaining a negative gap. The representative weighting comparison is in Supplementary Table S10, and AlphaFold pLDDT decomposed as a model score in Supplementary Table S9.

#### Robustness to dataset composition

We recompute the species-variance analysis after removing ribosomal proteins and on a domain-balanced subsample, using the same model representatives and predictor sets (Supplementary Section S3.4). We also recompute Elo without ribosomal families for ProteinMPNN, ESM-IF and ESM2-15B; numerical results and sample memberships are supplied in Source Data.

### Variance decomposition

For each model the standardised score is regressed by OLS on nested predictor sets (biophysics, family, species, and unions). We report the family-controlled increments Δ_bio_*_|_*_fam,spec_ and Δ_spec_*_|_*_fam,bio_ and their corresponding partial *R*^2^ values and nested *F* -tests, the species attenuation, and a within-family (Simpson’s) retention measure (Supplementary Section S3). All design matrices share one orthonormal basis so models are compared on identical predictor spaces.

To quantify how much of a model’s scoring behaviour is biophysics versus taxonomy, we decompose the variance of the score. Protein families are defined partly by enzyme class and GO terms, which co-vary with composition, and species differ systematically, so biophysics and taxonomy are confounded; the two are assessed jointly by comparing nested models rather than from any single regression.

#### Predictor sets

For each model the standardised score is regressed by OLS on each predictor set, and *R*^2^ = 1 *−* SSR*/*SST reported: LowDim (PC1+PC2 of the 14-feature PCA); Biophys (all 14 features); Family (*C*(protein_family)); Species (*C*(species)); the unions Family+Species, Biophys+Family, Biophys+Species; and Full (Biophys+Family+Species). The main variance comparisons use the 9,963 proteins with complete features and scores for all fourteen representatives. Each design matrix is factorised by column-pivoted QR, retaining its numerical rank, once and every model’s score projected onto the shared orthonormal basis, so all models are compared on identical predictor spaces.

### Family-controlled nested increments

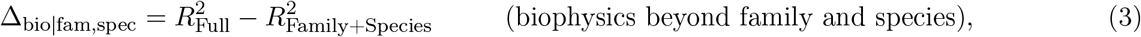

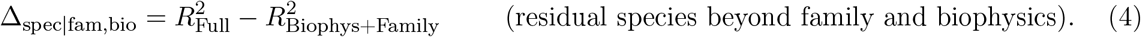

For each, the partial *R*^2^ = (SSR*_r_ −* SSR*_f_* )*/*SSR*_r_* and the nested *F* -test

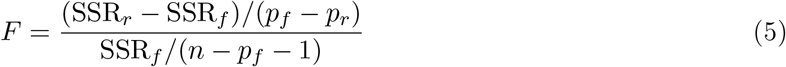

are reported. The categorical predictors carry many levels (species alone has 495), so their raw single-block *R*^2^ is dimensionality-inflated; interpretation rests on the increments, the attenuation, and the within-family retention, not on raw categorical *R*^2^. With large *n* every *F* -test is significant, so the relevant quantity is the effect size.

#### Total species effect and attenuation

The marginal effect is the single-block *R*_Species_^2^; the residual is Δ_spec_*_|_*_fam,bio_. Their relation gives the species attenuation,

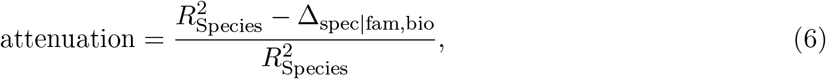

the fraction of species-associated score variance absorbed once family composition and biophysics are accounted for. Δ_spec_*_|_*_fam,bio_ is recomputed on three cohorts (full; ribosomal-removed; domain-balanced subsample) to check it is not a composition artefact.

#### Simpson’s-paradox check

A high marginal species effect can arise from a consistent within-family preference or from species differing in which families they contain. We compare reference-free per-species effects before and after removing family composition: the marginal effect is each species’ mean score; the adjusted effect is its mean after residualising on family (and family+biophysics). We report the count-weighted Pearson correlation, the retention = sd(adjusted)*/*sd(marginal), and the weighted fraction of species whose effect flips sign, all weighted by species *n*. Retention near 1 indicates a genuine within-family preference; collapse indicates the marginal signal was family composition.

### PCA and GAM landscapes

The 14 standardised features are decomposed by PCA (PC1+PC2 = 40.0 % of feature variance). Per model a GAM relates the z-scored score to (PC1, PC2) by REML (mgcv); its deviance explained is the preference variance, reported separately from feature variance. Grid cells with *<* 3 supporting proteins are masked. Per-domain overlap is quantified by Bhattacharyya overlap of the bivariate Gaussians.

### Design generation and analysis

Each model designs under one shared configuration (*T* = 0.1, 8 designs per template, full chain-A redesign). Designs and wild types are folded with ColabFold (single-sequence, 5 models / 3 recycles) ^64^; 16 features are computed on the rank-1 prediction; per-property paired shifts are reported as *d_z_*. Two-sided Wilcoxon signed-rank tests use a normal approximation with continuity correction, with paired differences rounded to 12 decimal places for ranking only. Benjamini–Hochberg FDR correction is applied jointly to all 84 tests across seven models and 12 varying properties. A per-feature mean-offset calibration places ColabFold designs in the AFDB PCA space without affecting within-predictor shift statistics. Functional-residue recovery uses curated UniProt annotations.

### Fine-tuning

ProteinMPNN (v_48_020; v_48_002 for sensitivity) was fine-tuned on alkaliphile- and acidophile-secretome cohorts, each matched 1:1 to neutralophile controls, with 40%-identity cluster-disjoint 70/15/15 splits and a validation-recovery guardrail. Held-out neutralophile backbones were redesigned and assessed using direct surface features. The surface acid–base PCA was fitted to four standardised surface features across 1,225 records (25 wild types and 200 designs from each of six primary or sensitivity models); positive PC1 was oriented toward higher surface acidic-residue fraction. Designs were refolded for self-consistency.

### Computational resources

The analysis was performed using CPU and GPU computing resources. The analysis reproduction workflow uses CPU resources.

## Data Availability

The protein-level dataset, including UniProt accession identifiers, is provided in Supplementary Data 1. Protein scores, biophysical features, design sequences, template-level observations and numerical figure and table data are supplied in the Source Data file. Links to the original UniProt and AlphaFold Database entries are included with the sequence metadata.

## Code Availability

The project repository, containing analysis code and reproduction instructions, is available at https://github.com/LBDillon/decoding-design-bias.

## Author Contributions Statement

L.B.D. performed the experiments and wrote the manuscript. O.M.C. supervised the project and contributed to the design of the experiments. A.M. contributed to building the fine-tuning component.

