## Supplementary Material for "Decoding the physicochemical basis of taxonomy preferences in protein design models"

**S11 Sequence-pretrained boundary cases** **33**

**S12 Tables supporting the main article** **34**

This Supplement follows the argument of the main manuscript. Sections S1–S2 define the evaluated models, analysis cohorts, scoring objects, generation protocols, and biophysical features. Sections S3–S5 quantify the magnitude and direction of taxonomic preference and test robustness to family composition, AlphaFold confidence, and structure source. Sections S6–S7 describe the biophysical organisation of model scores. Section S8 tests whether these preferences propagate into generated sequences. Sections S9–S10 examine their malleability under continued training and the structural compatibility of the resulting designs. Section S11 summarises models that occupy boundary positions between the principal scoring contexts.

All data and code supporting the findings are available in the project repository, with a scripted reproducer (`reproduce_optional/reproduce_key_results.sh`) and quick-start instructions.

### S1 Models, scoring objects, and sequence generation

#### S1.1 Model classes

The 14 scored models cover four mathematical scoring objects and three scoring contexts (Table S1)[1–11]. The principal distinctions are whether a supplied backbone is visible and whether the complete surrounding native sequence is visible when a residue is scored.

**Table S1:** Classes of protein design model evaluated in this study. Models are grouped by the information available when scoring each residue

| Scoring context | Models | Scored object | Target distribution |
|---|---|---|---|
| Backbone-conditioned | ProteinMPNN, SolubleMPNN, Caliby, SolubleCaliby, ESM-IF, TriFlow | autoregressive likelihood, energy, or predictive marginal | $p(\text{sequence} \mid \text{backbone})$ |
| Structure plus native-sequence context | MIF, MIF-ST, ESM3-structure | masked pseudo-log-likelihood | $p(s_i \mid \mathbf{s}_{\setminus i}, \mathbf{X})$ |
| Sequence-only | ESM2-15B, CARP-640M, ESM3-sequence, ProGen2-XL, ProtGPT2 | pseudo-log-likelihood or autoregressive likelihood | $p(\text{sequence})$ |

#### S1.2 Model specifications and training-corpus context

**Table S2:** Specifications of the 14 protein design models. Checkpoints and default hyperparameters were taken from the original repositories. Training-data descriptions are coarse because several published corpora combine multiple filtered sources.

| Model | Version/checkpoint | Architecture | Training data |
|---|---|---|---|
| ProteinMPNN | `v_48_020` | message-passing GNN | PDB structures with CATH-based splitting |
| SolubleMPNN | soluble `v_48_020` | message-passing GNN | soluble-protein PDB subset |
| Caliby | `afdb` | structure-conditioned Potts model | PDB-derived monomer structures |
| SolubleCaliby | soluble | structure-conditioned Potts model | soluble-protein structural subset |
| ESM-IF | `esm_if1_gvp4_t16_142M_UR50` | GVP-transformer | AlphaFold-DB models of UniRef sequences |
| TriFlow | `afdb_weights` | flow-matching model | AlphaFold-DB structures |
| ESM3-structure | ESM3-open | multimodal transformer | sequence and structure tokens |
| MIF | 1.0 | SE(3)-equivariant GNN | PDB/CATH structures |
| MIF-ST | 1.0 | GNN plus CARP-640M embeddings | PDB/CATH plus UniRef-derived sequence representations |
| ESM3-sequence | ESM3-open | multimodal transformer | sequence tokens with structure masked |
| ESM2-15B | 15B | transformer | UniRef50/90-derived sequence corpus |
| CARP-640M | 640M | dilated CNN | UniRef-derived sequence corpus |
| ProGen2-XL | XL checkpoint | autoregressive transformer | clustered protein-sequence databases |
| ProtGPT2 | published checkpoint | autoregressive transformer (BPE) | UniRef50 |

Experimentally determined structural corpora over-represent proteins that are expressible, soluble, ordered, stable, and experimentally tractable, while under-representing membrane proteins, flexible regions, low-complexity sequences, and intrinsically disordered proteins. Sequence corpora cover a broader taxonomic range but still reflect sequencing effort, annotation depth, reference-proteome selection, clustering procedures, and the over-representation of model organisms, pathogens, and medically or commercially important taxa. These distinct selection processes motivate separating preferences associated with scoring context from those associated with training-corpus composition.

##### S1.3 Likelihood scoring and cross-model comparability

Every model is used to score a fixed wild-type sequence in its available context and, where a stable generation procedure is available, to design sequences for a fixed backbone. Each score is reduced to a single per-residue quantity and oriented so that higher means more preferred: log-probabilities and negative energies are retained, while negative log-probabilities and energies are sign-flipped. Positions with non-standard residues or missing coordinates are excluded from both the numerator and denominator.

**Autoregressive conditional log-likelihood.**
ProteinMPNN, SolubleMPNN, AlkSecMPNN, AcidSecMPNN, and ESM-IF are structure-conditioned; ProGen2-XL and ProtGPT2 are sequence-only. Given a decoding order $\pi$,

$$\ell_{\mathrm{AR}}(\mathbf{s} \mid \mathbf{X}) = \frac{1}{L}\sum_{i=1}^{L} \log p\big(s_{\pi(i)} \mid s_{\pi(<i)}, \mathbf{X}\big)\,, \tag{S1}$$

where the scored residue is hidden. ProteinMPNN-family models are scored with `protein_mpnn_run.py -score_only 1`; we store $-$`global_score`. ESM-IF uses the per-residue mean `ll_fullseq`. Sequence-only autoregressive models omit $\mathbf{X}$ and use the natural left-to-right order. ProtGPT2 uses byte-pair encoding, so its per-token likelihood is normalised by residue count and token count is retained as a covariate.

**Masked-marginal pseudo-log-likelihood.** MIF and MIF-ST use sequence plus backbone; ESM2-15B, CARP-640M, and ESM3-sequence use sequence alone; ESM3-structure uses structure plus visible surrounding native sequence. Each position is masked in turn:

$$\mathrm{PLL}(\mathbf{s} \mid [\mathbf{X}]) = \frac{1}{L}\sum_{i=1}^{L} \log p\big(s_i \mid \mathbf{s}_{\setminus i}, [\mathbf{X}]\big)\,. \tag{S2}$$

ESM3 is run in two modes differing only in whether structure tokens are supplied. In both modes, all surrounding native residues remain visible; ESM3-structure is therefore classified as structure plus native-sequence context rather than backbone-only inverse folding.

**Flow-matching predictive marginal.** TriFlow integrates one Euler denoising trajectory from an all-mask sequence while conditioning on the clean backbone (`noise_label`=0, `temp`=1.0, no guidance) and accumulates the wild-type residue probability, $L^{-1}\sum_i \log p(s_i \mid \mathbf{X})$. The score is stochastic.

**Potts pseudo-energy.** Caliby and SolubleCaliby fit a Potts model to the input structure and evaluate

$$U(\mathbf{s} \mid \mathbf{X}) = -\sum_{i} h_i(s_i) - \sum_{i<j} J_{ij}(s_i, s_j), \tag{S3}$$

with $-U/L$ stored as the oriented score.

**Valid cross-model comparisons.** These families define different mathematical objects with different visibility, normalisation, and tokenisation. Because of this we do not compare absolute score magnitudes across models. Comparisons are restricted to per-model variance decomposition, within-model ranks and residuals, explained-variance patterns, and model-normalised contrasts. Species Elo z-scores each model within protein family before forming pairwise matches. Property-importance analyses residualise each score on a unigram-composition plus length baseline where specified.

##### S1.4 Sequence-generation protocol

Seven models generated designs for the 25 templates in Section S8.1: ProteinMPNN, SolubleMPNN, Caliby, SolubleCaliby, ESM-IF, MIF, and MIF-ST. Generation used a shared configuration where supported: sampling temperature $T = 0.1$; eight sequences per template (seeds 0–7); temperature-only sampling (top-$k = 0$, top-$p = 1$); no omitted amino acids; full redesign of chain A; and the same AlphaFold-DB v6 monomer as structural input[12]. Output tables were accepted only after checks for canonical residues, WT-matched length, expected design count, temperature consistency, and non-trivial divergence from WT. MIF's nucleus filters were disabled; ESM-IF used `GVPTransformerModel.sample`; MIF and MIF-ST iteratively unmasked a fully masked sequence; and Caliby-family models Gibbs-sampled the fitted Potts distribution.

#### S2 Analysis cohorts and biophysical features

##### S2.1 Analysis cohorts

The study uses distinct cohorts for scoring, redesign, structural robustness, and fine-tuning (Table S3). Keeping these cohorts explicit prevents length ranges, structure sources, and sample sizes from being conflated.

**Table S3:** Overview of the principal analysis cohorts. Pair counts for the extremophile cohorts are approximate because final counts depend on quality-control and split filters.

| Cohort | Size | Structural input | Primary purpose |
|---|---|---|---|
| Main natural-protein cohort | 10,146 proteins; 495 species; 281 families | AlphaFold-DB v6 | likelihood scoring, variance decomposition, Elo, PCA/GAM, and property importance |
| Design-template cohort | 25 templates; 8 designs per model | AlphaFold-DB v6; ColabFold refolds | WT-to-design property shifts and functional-residue recovery |
| Experimental-PDB cohort | 876 chains; 213 species; 185 families | X-ray structures at $\leq$ 2.5 Å | robustness to structure source and resolved-chain sequence input |
| Alkaliphile secretome pairs | about 250 matched pairs | AlphaFold-derived backbones | ProteinMPNN continued training and held-out surface redesign |
| Acidophile secretome pairs | about 75 matched pairs | AlphaFold-derived backbones | opposite-direction ProteinMPNN continued training |
| Matched ESM2 comparison | 10 held-out secreted targets | same exposed positions; backbone used only by ProteinMPNN | sequence- versus structure-conditioned malleability |

The main scoring cohort spans 24–2531 amino acids and is not length-filtered. The redesign cohort spans 49–633 amino acids and was selected for tractable folding and generation while covering all three domains, multiple structural classes, and the full range of ProteinMPNN Elo preference classes.

#### S2.2 Biophysical feature definitions

Sixteen global properties are computed per protein, either deterministically from amino-acid sequence using BioPython `ProteinAnalysis`[13] or from the AlphaFold backbone using `paper_code/shared/features/`. Fourteen features form the common modelling set used for PCA, variance decomposition, and property importance: nine sequence-derived and five structure-derived features. Net charge at pH 7 is excluded because it is collinear with pI ($|r| = 0.85$), and small-residue fraction is excluded because it is collinear with molecular weight per residue ($|r| = 0.86$). The maximum pairwise correlation among retained features is $|r| = 0.74$. Fine-tuning analyses additionally compute surface-specific charge quantities, including surface net charge and charge per residue; these are not part of the 14-feature global PCA.

**Table S4:** Sequence-derived properties. Nine enter the common 14-feature modelling set. The two italicised properties are computed for supplementary read-outs but excluded from the common set owing to collinearity. Residue sets are acidic = $\{D, E\}$, basic = $\{K, R, H\}$, aromatic = $\{F, W, Y\}$, and small = $\{A, G, S, T\}$.

| Feature | Definition | Interpretation |
|---|---|---|
| Sequence length | Total number of amino acids | Longer means a larger protein |
| MW per residue | Molecular weight divided by sequence length | Higher values indicate heavier residue composition |
| Isoelectric point (pI) | pH at which predicted net charge is zero | Values below 7 are more acidic; above 7 more basic |
| *Net charge at pH 7* | Predicted net charge at physiological pH | Positive means an excess of basic residues |
| Acidic-residue fraction | Fraction of Asp and Glu | Higher means more acidic composition |
| Basic-residue fraction | Fraction of Lys, Arg, and His | Higher means more basic composition |
| GRAVY | Grand average of Kyte–Doolittle hydropathy | Negative is hydrophilic; positive hydrophobic |
| Aromaticity | Fraction of Phe, Trp, and Tyr | Higher means more aromatic composition |
| Instability index | Dipeptide-frequency stability predictor | Values above 40 predict lower stability |
| Proline fraction | Fraction of Pro residues | Higher means more proline and backbone rigidity |
| *Small-residue fraction* | Fraction of Ala, Gly, Ser, and Thr | Higher means more small or flexible residues |

**Table S5:** Structure-derived features in the common 14-feature modelling set.

| Feature | Definition | Interpretation |
|---|---|---|
| Relative contact order (RCO) | Mean sequence separation of contacting residues ($C\alpha$–$C\alpha \leq 8$ Å), normalised by length | Higher means more long-range topology |
| Mean $C\beta$ distance | Mean distance of $C\beta$ atoms ($C\alpha$ for Gly) from the centroid | Higher means a more extended structure |
| Surface exposure | Fraction of residues beyond the 70th percentile of $C\alpha$–centroid distances | Higher means a larger exposed fraction |
| Helix–sheet contrast | Helix percentage minus sheet percentage | Positive means helix-dominant |
| Ordered percentage | Helix percentage plus sheet percentage | Higher means more regular secondary structure |

### S3 Variance decomposition and within-family robustness

#### S3.1 Nested variance-decomposition framework

To determine how much apparent taxonomic preference survives adjustment for protein type and measured biophysics, we decompose the variance of each model's score using nested ordinary-least-squares models. Biophysics, family, and species are confounded, so interpretation rests on increments between nested models rather than a single raw regression.

For each model, the standardised score is regressed on the following predictor sets:

- LowDim: PC1 and PC2 of the 14-feature PCA;
- Biophys: all 14 standardised features;
- Family: $C(\texttt{protein\_family})$;
- Species: $C(\texttt{species})$;
- all pairwise unions of Biophys, Family, and Species;
- Full: Biophys plus Family plus Species.

Each design matrix is QR-factorised once and each model's score is projected onto the same orthonormal predictor basis. The low-dimensional term measures the fraction of score variance captured linearly by the displayed PCA plane; GAM deviance explained is reported separately in Section S6.3.

The family-controlled increments are

$$\Delta_{\text{bio}|\text{fam,spec}} = R^2_{\text{Full}} - R^2_{\text{Family+Species}}, \tag{S4}$$

$$\Delta_{\text{spec}|\text{fam,bio}} = R^2_{\text{Full}} - R^2_{\text{Biophys+Family}}. \tag{S5}$$

For each comparison, the partial $R^2$ is

$$R^2_{\text{partial}} = \frac{\text{SSR}_r - \text{SSR}_f}{\text{SSR}_r}, \tag{S6}$$

where $r$ and $f$ denote reduced and full models. The nested $F$ statistic is

$$F = \frac{(\text{SSR}_r - \text{SSR}_f)/(p_f - p_r)}{\text{SSR}_f/(n - p_f - 1)}. \tag{S7}$$

At $n \approx 10{,}000$, all tests are statistically significant; interpretation therefore rests on effect size.

#### S3.2 Species attenuation and within-family retention

The marginal species effect is the single-block $R^2_{\text{Species}}$. Species attenuation is

$$\text{attenuation} = \frac{R^2_{\text{Species}} - \Delta_{\text{spec}|\text{fam,bio}}}{R^2_{\text{Species}}}, \tag{S8}$$

which measures the fraction of species-associated score variance absorbed once protein-family composition and measured biophysics are included.

A separate within-family check distinguishes a consistent taxonomic preference from a change in the families represented by each species. The marginal species effect is each species' mean standardised score. The adjusted effect is its mean score after residualising on family, and then on family plus biophysics. We report the count-weighted correlation between marginal and adjusted effects, the retention

$$\text{retention} = \frac{\text{sd(adjusted species effects)}}{\text{sd(marginal species effects)}}, \tag{S9}$$

and the weighted fraction of species whose effect changes sign. Effects with absolute magnitude at most $10^{-12}$ in standardised-score units are treated as numerical zero and do not define a direction reversal. Retention near one indicates that the species ordering survives within families; retention near zero indicates that the marginal taxonomic signal arose mainly from family composition.

##### S3.3 Main decomposition results

Measured biophysics explains substantial unique variance in backbone-conditioned scores but much less in classical sequence-only language-model scores. The residual species increment shows the opposite pattern. Backbone-conditioned models attenuate 91–93% of their apparent species signal and retain only 0.22–0.28 of the marginal per-species effect within families. ESM2-15B, CARP-640M, ProGen2-XL, and ProtGPT2 attenuate only 15–29% and retain 0.76–0.82. MIF-ST is sequence-like despite receiving a backbone; ESM3-sequence is a boundary case with low residual species dependence despite being scored without supplied structure.

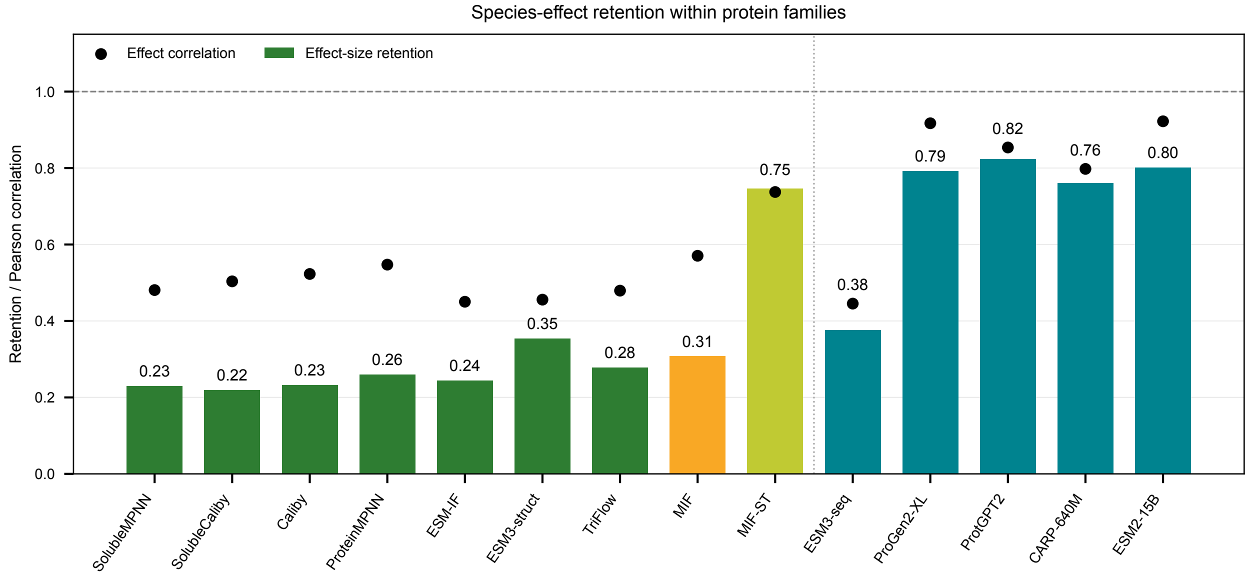

**Figure S1:** Within-family retention of species effects across 14 models. Coloured bars show the ratio of the standard deviation of adjusted species effects to that of marginal species effects; black points show their Pearson correlation. Adjustment includes family and the 14 biophysical features. Dark green, amber, lime and teal denote backbone-conditioned, MIF, MIF-ST and sequence-only models, respectively; ESM3-structure is grouped with structurally conditioned models for this display. Each bar or point is one derived model-level estimate from the common complete-case cohort ($n = 9{,}963$ proteins); no error bars are shown. Source data are provided as a Source Data file.

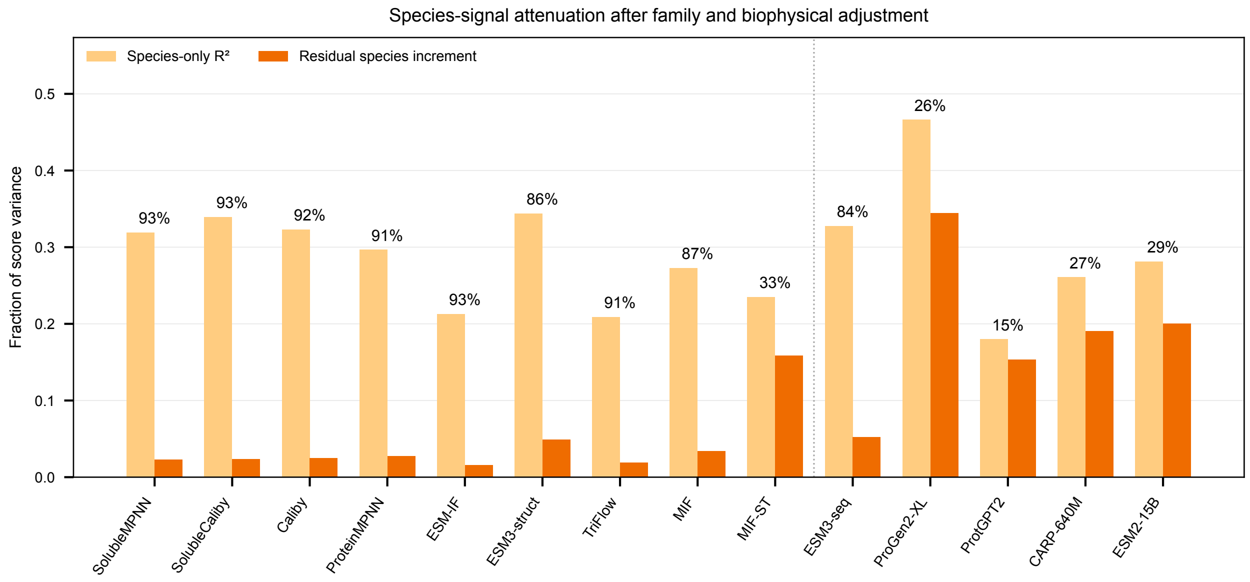

**Figure S2:** Apparent species signal before and after adjustment. Pale-orange bars show species-only $R^2$; dark-orange bars show the residual species increment after including family and biophysics. Labels report the percentage attenuation. Each bar is one derived model estimate from the common complete-case cohort ($n = 9{,}963$ proteins), rather than a replicate mean. Source data are provided as a Source Data file.

**Table S6:** Score-variance decomposition per model (complete-case $n = 9{,}963$). Biophys $R^2$ is the variance explained by the 14 features. Biophysics partial $R^2$ is the increment beyond family plus species, reported as partial $R^2$. Marginal species $R^2$ is the species block alone. $\Delta_{\text{spec}}$ is residual species variance beyond family plus biophysics, expressed as a fraction of total score variance. All nested $F$ tests have $p < 10^{-3}$.

| Model | Context | Biophys $R^2$ (%) | Bio. partial $R^2$ | Marg. species $R^2$ (%) | $\Delta_{\text{spec}}$ (%) | Attenuation (%) | Retention |
|---|---|---|---|---|---|---|---|
| ProteinMPNN | backbone | 64.3 | 0.19 | 29.7 | 2.8 | 91 | 0.26 |
| SolubleMPNN | backbone | 66.9 | 0.18 | 31.9 | 2.3 | 93 | 0.23 |
| Caliby | backbone | 65.4 | 0.21 | 32.3 | 2.5 | 92 | 0.23 |
| SolubleCaliby | backbone | 66.6 | 0.21 | 33.9 | 2.3 | 93 | 0.22 |
| ESM-IF | backbone | 47.2 | 0.34 | 21.3 | 1.6 | 93 | 0.24 |
| TriFlow | backbone | 46.9 | 0.11 | 20.9 | 1.9 | 91 | 0.28 |
| MIF | structure+sequence | 57.3 | 0.15 | 27.3 | 3.4 | 87 | 0.31 |
| MIF-ST | structure+sequence | 27.8 | 0.07 | 23.5 | 15.9 | 33 | 0.75 |
| ESM3-structure | structure+sequence | 47.5 | 0.13 | 34.4 | 4.9 | 86 | 0.35 |
| ESM3-sequence | sequence-only | 42.8 | 0.11 | 32.8 | 5.2 | 84 | 0.38 |
| ESM2-15B | sequence-only | 7.3 | 0.03 | 28.1 | 20.1 | 29 | 0.80 |
| CARP-640M | sequence-only | 18.2 | 0.05 | 26.1 | 19.0 | 27 | 0.76 |
| ProGen2-XL | sequence-only | 14.3 | 0.04 | 46.6 | 34.5 | 26 | 0.79 |
| ProtGPT2 | sequence-only | 22.9 | 0.04 | 18.0 | 15.3 | 15 | 0.82 |

#### S3.4 Robustness across dataset composition

The residual species increment was recomputed on the common complete-case cohort ($n = 9{,}963$), after removing ribosomal proteins ($n = 6{,}779$), and on a domain-balanced subsample ($n = 4{,}059$; 1,353 proteins per domain). The subsample was drawn without replacement with R seed 0; its accession membership is supplied in Source Data. All three analyses retain the same fourteen representatives, and the same fourteen biophysical features. The separation between backbone-conditioned and classical sequence-only language models persists. These checks address the effects of the over-represented ribosomal class and marginal domain proportions.

### S4 Robustness to structural-input artefacts

#### S4.1 Independent experimental-structure cohort

To test the main result on experimentally determined structures, we assembled an independent RCSB PDB cohort rather than selecting the small experimentally solved subset of the main dataset. The Search API (v2) was queried for X-ray protein entities at resolution $\leq 2.5$ Å, sequence length 50–1000, from Bacteria, Eukaryota, or Archaea, and restricted to single-chain monomeric assemblies. The search returned 67,812 entities, collapsed to 10,918 representatives at 30% sequence identity. After Data API and UniProt annotation, taxonomic filtering, family assignment, and the same phylogenetic-breadth criteria as the main cohort, the set contained 2,776 proteins. A fixed-seed subsample matched the main cohort's domain marginal, giving 876 chains from 213 species and 185 families at median resolution 1.95 Å.

For each representative, the resolved chain was extracted and its resolved-coordinate sequence was scored. Eighty-five per cent of resolved chains are truncated relative to the deposited construct and no chain is identical to the complete UniProt sequence, so all models, including sequence-only models, receive inputs that differ from the main analysis. The cohort is nearly ribosomal-free (1.5% versus 31.4% in the main cohort), reflecting the scarcity of single-chain monomeric ribosomal X-ray structures.

#### S4.2 Matched AlphaFold-DB control

A direct comparison with the full AlphaFold-DB cohort confounds structure source with cohort size, composition and model-specific score coverage. We fitted the same 14-feature, family and species predictor blocks using column-pivoted QR, retaining only the numerical column space. Numerical rank, including the intercept, determines the residual degrees of freedom and adjusted $R^2$. This removes redundant family/species columns; an unpivoted QR with all returned columns can fit directions outside the predictor space. Residual species variance is the increment in $R^2$ after adding species to biophysics and family; its adjusted counterpart is the difference between the two adjusted $R^2$ values.

PDB fits use each model's complete cases. For each model, we prefiltered the AlphaFold-DB pool for available scores, features and group labels, excluded ribosomal proteins, and drew 30 samples without replacement (seeds 1000–1029), matching the exact size and taxonomic-domain counts of that model's scored PDB subset. Thus the matched sample sizes range from 562 to 876 rather than assuming that every model scored all 876 chains. All draw memberships and model-specific counts are supplied in Source Data. The same score column is used on both structure sources; ProGen2-XL uses `progen2_XL_score` in this control.

We also repeated the analysis on the 551 PDB proteins scored by all twelve models, with controls matched to that shared subset (Eukaryota 231, Bacteria 247, Archaea 73). Fits and degrees of freedom were independently verified in R. Control dispersions are descriptive standard deviations across draws, not confidence intervals for a causal structure-source effect.

**Table S7:** AlphaFold-DB comparison input and experimental-PDB cohort universe. Model-specific complete cases and exactly size/domain-matched AFDB controls are used in the analyses below.

| Property | AFDB comparison input | Experimental-PDB cohort |
|---|---|---|
| Proteins | 10,146 | 876 |
| Structure source | AlphaFold2 model | experimental X-ray |
| Sequence scored | complete UniProt | resolved chain |
| Eukaryota (%) | 45.0 | 45.2 |
| Bacteria (%) | 41.6 | 41.6 |
| Archaea (%) | 13.4 | 13.2 |
| Distinct families | 281 | 185 |
| Distinct species | 495 | 213 |
| Ribosomal (%) | 31.4 | 1.5 |
| Median length (aa) | 259 | 269 |
| Median resolution (Å) | – | 1.95 |

#### S4.3 Experimental-cohort results

Raw residual-species estimates show positive agreement across the twelve retained models between PDB and the means of their size/domain-matched AFDB controls (Spearman $\rho = 0.762$, nominal $p = 0.004$; Pearson $r = 0.811$). ProteinMPNN has a raw increment of 0.124 on 875 PDB chains versus $0.032 \pm 0.004$ in its matched controls; ESM2-15B gives 0.264 versus $0.265 \pm 0.025$, and ProtGPT2 0.133 versus $0.133 \pm 0.011$ (Supplementary Table S8). PDB estimates for backbone-conditioned models range from 0.124 to 0.171, overlapping the classical sequence-model range of 0.133 to 0.276. There is therefore no uniform ordering of these groups in the experimental cohort.

Agreement is sensitive to the statistic and cohort. Comparing adjusted increments between PDB and the full AFDB input gives Spearman $\rho = 0.420$ (nominal $p = 0.175$), despite Pearson $r = 0.807$. On the shared 551-protein PDB subset, raw increments correlate with matched-control means (Spearman $\rho = 0.678$, nominal $p = 0.015$), but the adjusted increments do not retain that ordering ($\rho = -0.084$, nominal $p = 0.795$). These correlations describe related, selected model outputs; their nominal $p$ values do not represent independent biological replication. The control provides partial agreement in raw score patterns, with substantial sensitivity to score coverage, cohort and degrees-of-freedom adjustment. It does not establish a structure-source-invariant ranking of models.

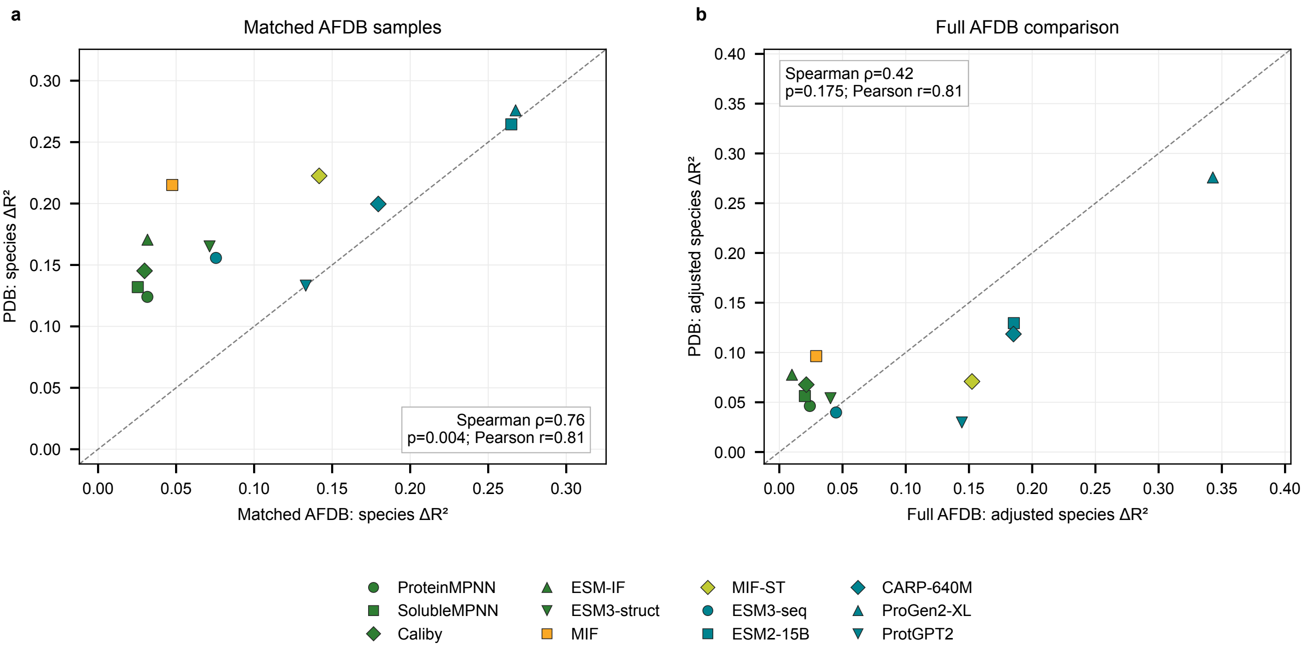

**Figure S3:** Rank-corrected experimental-PDB robustness comparison across twelve model outputs. a, Raw residual-species $R^2$ increments on PDB versus means of 30 AFDB draws matched to each model's actual PDB sample size and domain counts (562–876 proteins). b, Adjusted-$R^2$ increments on PDB versus the same-score full AFDB input. Dashed lines indicate equality. Model names are identified by the colour and symbol key below the panels. Green denotes structural input, orange MIF, light green MIF-ST and teal sequence-mode models; distinct symbols separate models within each colour group. Points are derived model estimates, not replicate means; control standard deviations are in Supplementary Table S8. Correlations are descriptive across related models and their $p$ values are nominal. SolubleCaliby is omitted because its raw PDB score export duplicates Caliby's. All input rows, draw memberships, rank diagnostics and the shared-complete-case sensitivity are supplied. Source data are provided as a Source Data file.

**Table S8:** Rank-corrected residual species variance and within-family retention on PDB and matched AFDB controls. Each model uses $n$ scored PDB proteins and 30 AFDB draws with exactly the same $n$ and domain counts. Δspec is the raw $R^2$ increment; adj. is the adjusted-$R^2$ increment; ret. is the ratio of adjusted to marginal species-effect standard deviations. AFDB values are draw means and s.d. is the population standard deviation across those 30 draws. Negative adjusted increments, when present, are retained. SolubleCaliby is omitted because its raw PDB score export duplicates Caliby's. Full observations and shared-cohort sensitivity are in Source Data.

| Model | $n$ | Δspec PDB | Δspec AFDB | AFDB s.d. | adj. PDB | adj. AFDB | ret. PDB | ret. AFDB |
|---|---|---|---|---|---|---|---|---|
| ProteinMPNN | 875 | 0.124 | 0.032 | 0.004 | 0.046 | 0.018 | 0.472 | 0.212 |
| SolubleMPNN | 875 | 0.132 | 0.025 | 0.003 | 0.056 | 0.015 | 0.489 | 0.189 |
| Caliby | 870 | 0.145 | 0.030 | 0.004 | 0.068 | 0.017 | 0.541 | 0.204 |
| ESM-IF | 847 | 0.171 | 0.032 | 0.004 | 0.078 | 0.007 | 0.562 | 0.234 |
| ESM3-structure | 876 | 0.165 | 0.071 | 0.007 | 0.054 | 0.034 | 0.571 | 0.355 |
| MIF | 562 | 0.215 | 0.048 | 0.006 | 0.096 | 0.024 | 0.568 | 0.241 |
| MIF-ST | 562 | 0.223 | 0.142 | 0.018 | 0.071 | 0.103 | 0.543 | 0.471 |
| ESM3-sequence | 876 | 0.156 | 0.076 | 0.008 | 0.040 | 0.045 | 0.561 | 0.371 |
| ESM2-15B | 876 | 0.264 | 0.265 | 0.025 | 0.129 | 0.181 | 0.679 | 0.660 |
| CARP-640M | 876 | 0.200 | 0.180 | 0.014 | 0.119 | 0.149 | 0.513 | 0.578 |
| ProGen2-XL | 876 | 0.276 | 0.268 | 0.023 | 0.276 | 0.270 | 0.523 | 0.574 |
| ProtGPT2 | 876 | 0.133 | 0.133 | 0.011 | 0.030 | 0.101 | 0.475 | 0.519 |

The PDB analysis tests consistency rather than isolating structure source. Selection favours rigid, well-ordered monomers; the archaeal subset contains only 17 mostly hyperthermophilic species; and resolved chains omit unmodelled residues. A direct structure-source comparison would require scoring predicted and experimental structures of the same proteins using identical sequences, which was not available here.

##### S4.4 AlphaFold confidence is largely family-driven

We treated average AlphaFold pLDDT as an additional response and applied the same 14-feature variance decomposition on the full cohort. Family alone explains 74% of pLDDT variance and the 14 biophysical features explain 38%. The residual species term after family and biophysics is 1.6% (0.7% adjusted), with 90% attenuation and within-family retention 0.30. pLDDT therefore shows the same family-driven signature as backbone-conditioned scores and a much smaller residual species term than classical sequence-only language models.

**Table S9:** AlphaFold confidence decomposed as a model response on the full cohort, beside representative scoring models.

| Response | Context | Biophys $R^2$ | residual species | attenuation | retention |
|---|---|---|---|---|---|
| AlphaFold pLDDT | structural confidence | 0.38 | 0.016 | 90% | 0.30 |
| ProteinMPNN | backbone | 0.64 | 0.028 | 91% | 0.26 |
| ESM-IF | backbone | 0.47 | 0.016 | 93% | 0.24 |
| MIF | structure+sequence | 0.57 | 0.034 | 87% | 0.31 |
| ESM2-15B | sequence-only | 0.07 | 0.201 | 29% | 0.80 |
| CARP-640M | sequence-only | 0.18 | 0.190 | 27% | 0.76 |
| ProGen2-XL | sequence-only | 0.14 | 0.345 | 26% | 0.79 |

#### S5 Taxonomic preference direction and Elo robustness

##### S5.1 Species-level Elo

Taxonomic preference direction is quantified per model with a species-level Elo rating following Ding and Steinhardt[14,15]. Within each protein family, model scores are z-scored and pairwise comparisons between species update ratings according to

$$R'_A = R_A + K(S - E_A), \qquad E_A = \frac{1}{1 + 10^{(R_B - R_A)/400}}, \tag{S10}$$

with $K = 32$, baseline 1500, and averaging over 50 random match orderings. Ratings cover all 495 species overall (492 for ESM2-15B, owing to unavailable scores).

The unweighted score Elo is primary. Two sensitivity variants are also computed: a pLDDT-weighted Elo and a pLDDT-residual Elo, in which each model's score is first regressed on average pLDDT and the Elo is recomputed on residuals. For ProteinMPNN, ESM-IF and ESM2-15B, unweighted and weighted ratings agree almost perfectly (Spearman $\rho$ = 0.998–0.999), with identical top-rated domains. Residualisation attenuates the Archaea–Eukaryota gap but does not reverse the gap for ProteinMPNN or ESM-IF.

**Table S10:** Archaea–Eukaryota Elo gaps under the primary unweighted rating and pLDDT-residual control. The pLDDT-weighted gaps are +468 for ProteinMPNN, +214 for ESM-IF and −99 for ESM2-15B; the corresponding species-level ratings are supplied in Source Data.

| Model | Context | Unweighted | pLDDT-residual |
|---|---|---|---|
| ProteinMPNN | backbone | +496 | +427 |
| ESM-IF | backbone | +229 | +110 |
| ESM2-15B | sequence-only | −98 | −104 |

##### S5.2 Full domain-level Elo statistics

Most backbone-conditioned models favour Archaea or thermophilic Bacteria, whereas the classical sequence language models CARP-640M, ProGen2-XL, and ProtGPT2 favour Eukaryota. MIF and MIF-ST provide an internal contrast: MIF favours Archaea, while addition of CARP-derived sequence representations in MIF-ST reverses the direction toward Eukaryota.

**Table S11:** Per-domain species Elo under the primary unweighted rating. Values are means across species; SEM equals SD divided by the square root of the domain species count.

| Model (context) | Domain | Mean | SD | SEM |
|---|---|---|---|---|
| ProteinMPNN (backbone) | Archaea | 1800.1 | 287.9 | 34.9 |
| | Bacteria | 1565.1 | 214.8 | 13.8 |
| | Eukaryota | 1304.6 | 203.2 | 14.9 |
| SolubleMPNN (backbone) | Archaea | 1799.9 | 274.8 | 33.3 |
| | Bacteria | 1559.4 | 228.6 | 14.7 |
| | Eukaryota | 1312.0 | 220.8 | 16.2 |
| Caliby (backbone) | Archaea | 1769.7 | 261.1 | 31.7 |
| | Bacteria | 1571.8 | 216.5 | 13.9 |
| | Eukaryota | 1306.9 | 211.0 | 15.5 |
| SolubleCaliby (backbone) | Archaea | 1782.8 | 263.7 | 32.0 |
| | Bacteria | 1581.7 | 221.2 | 14.2 |
| | Eukaryota | 1289.2 | 217.9 | 16.0 |
| ESM-IF (backbone) | Archaea | 1581.4 | 237.1 | 28.7 |
| | Bacteria | 1590.2 | 223.6 | 14.4 |
| | Eukaryota | 1352.0 | 201.0 | 14.8 |
| TriFlow (backbone) | Archaea | 1660.1 | 213.8 | 25.9 |
| | Bacteria | 1541.4 | 207.8 | 13.4 |
| | Eukaryota | 1387.0 | 174.5 | 12.8 |
| MIF (structure+sequence) | Archaea | 1730.8 | 294.8 | 35.7 |
| | Bacteria | 1573.4 | 212.3 | 13.6 |
| | Eukaryota | 1319.1 | 191.3 | 14.1 |
| MIF-ST (structure+sequence) | Archaea | 1277.7 | 218.0 | 26.4 |
| | Bacteria | 1474.2 | 232.9 | 15.0 |
| | Eukaryota | 1615.5 | 277.3 | 20.4 |
| ESM3-structure (structure+sequence) | Archaea | 1415.0 | 229.6 | 27.8 |
| | Bacteria | 1590.3 | 256.5 | 16.5 |
| | Eukaryota | 1413.2 | 238.9 | 17.6 |
| ESM3-sequence (sequence-only) | Archaea | 1357.5 | 244.0 | 29.6 |
| | Bacteria | 1590.5 | 269.6 | 17.3 |
| | Eukaryota | 1434.0 | 263.1 | 19.3 |
| ESM2-15B (sequence-only) | Archaea | 1398.6 | 219.2 | 26.6 |
| | Bacteria | 1531.2 | 244.9 | 15.8 |
| | Eukaryota | 1497.0 | 264.2 | 19.4 |
| CARP-640M (sequence-only) | Archaea | 1154.1 | 217.1 | 26.3 |
| | Bacteria | 1460.4 | 234.3 | 15.1 |
| | Eukaryota | 1678.9 | 288.4 | 21.2 |
| ProGen2-XL (sequence-only) | Archaea | 1065.9 | 269.3 | 32.7 |
| | Bacteria | 1503.6 | 320.2 | 20.6 |
| | Eukaryota | 1654.8 | 411.3 | 30.2 |
| ProtGPT2 (sequence-only) | Archaea | 1131.4 | 208.3 | 25.3 |
| | Bacteria | 1440.9 | 215.4 | 13.8 |
| | Eukaryota | 1712.8 | 240.9 | 17.7 |

**Table S12:** Domain-level ANOVA and pairwise Cohen's $d$ for species Elo. All ANOVA tests have $p < 10^{-3}$. Effect-size signs follow the first named domain.

| Model | Context | ANOVA $F$ | Bac–Arch | Bac–Euk | Arch–Euk |
|---|---|---|---|---|---|
| ProteinMPNN | backbone | 144.1 | −1.01 | +1.24 | +2.17 |
| SolubleMPNN | backbone | 124.8 | −1.00 | +1.10 | +2.06 |
| Caliby | backbone | 133.9 | −0.87 | +1.24 | +2.05 |
| SolubleCaliby | backbone | 149.2 | −0.87 | +1.33 | +2.14 |
| ESM-IF | backbone | 68.5 | +0.04 | +1.11 | +1.09 |
| TriFlow | backbone | 58.3 | −0.57 | +0.80 | +1.47 |
| MIF | structure+sequence | 115.3 | −0.68 | +1.25 | +1.84 |
| MIF-ST | structure+sequence | 48.4 | +0.86 | −0.56 | −1.29 |
| ESM3-structure | structure+sequence | 31.8 | +0.70 | +0.71 | +0.01 |
| ESM3-sequence | sequence-only | 29.9 | +0.88 | +0.59 | −0.30 |
| ESM2-15B | sequence-only | 7.5 | +0.55 | +0.13 | −0.39 |
| CARP-640M | sequence-only | 112.1 | +1.33 | −0.84 | −1.94 |
| ProGen2-XL | sequence-only | 69.9 | +1.41 | −0.42 | −1.56 |
| ProtGPT2 | sequence-only | 183.4 | +1.45 | −1.20 | −2.50 |

**Table S13:** Representative top-ranked species patterns under the unweighted Elo. Interpretations refer to reported training-corpus composition and should be read as descriptive rather than causal.

| Training data/modality | Model | Representative species-level preference |
|---|---|---|
| Structure-only | ProteinMPNN | Archaeal hyperthermophiles (*Pyrococcus*, *Thermococcus*) and *Thermus thermophilus*. |
| | ESM-IF | Thermophilic and extremophile bacteria, including *Caldanaerobacter*, *Geobacillus*, and *Thermus*. |
| | Caliby | Archaeal and thermophilic extremophiles. |
| | TriFlow | Thermophilic bacteria and archaea. |
| CATH structure+sequence | MIF | Archaeal thermophiles; nine of its ten highest-ranked species are archaeal. |
| CATH plus UniRef representations | MIF-ST | Mammalian and domesticated eukaryotes, including sheep, cattle, pig, and dog. |
| ESM family | ESM3-structure | Pathogenic and thermophilic bacteria. |
| | ESM3-sequence | Enterobacterial and thermophilic bacteria. |
| | ESM2-15B | Enterobacterial pathogens, including *Salmonella*, *Yersinia*, and *E. coli*. |
| Sequence-only | CARP-640M | Vertebrate model organisms; ten of ten top species are eukaryotic. |
| | ProGen2-XL | Predominantly plant and animal species; nine of its ten highest-ranked species are eukaryotic. |
| | ProtGPT2 | Vertebrates, including dog, sheep, cat, and cattle; ten of ten top species are eukaryotic. |

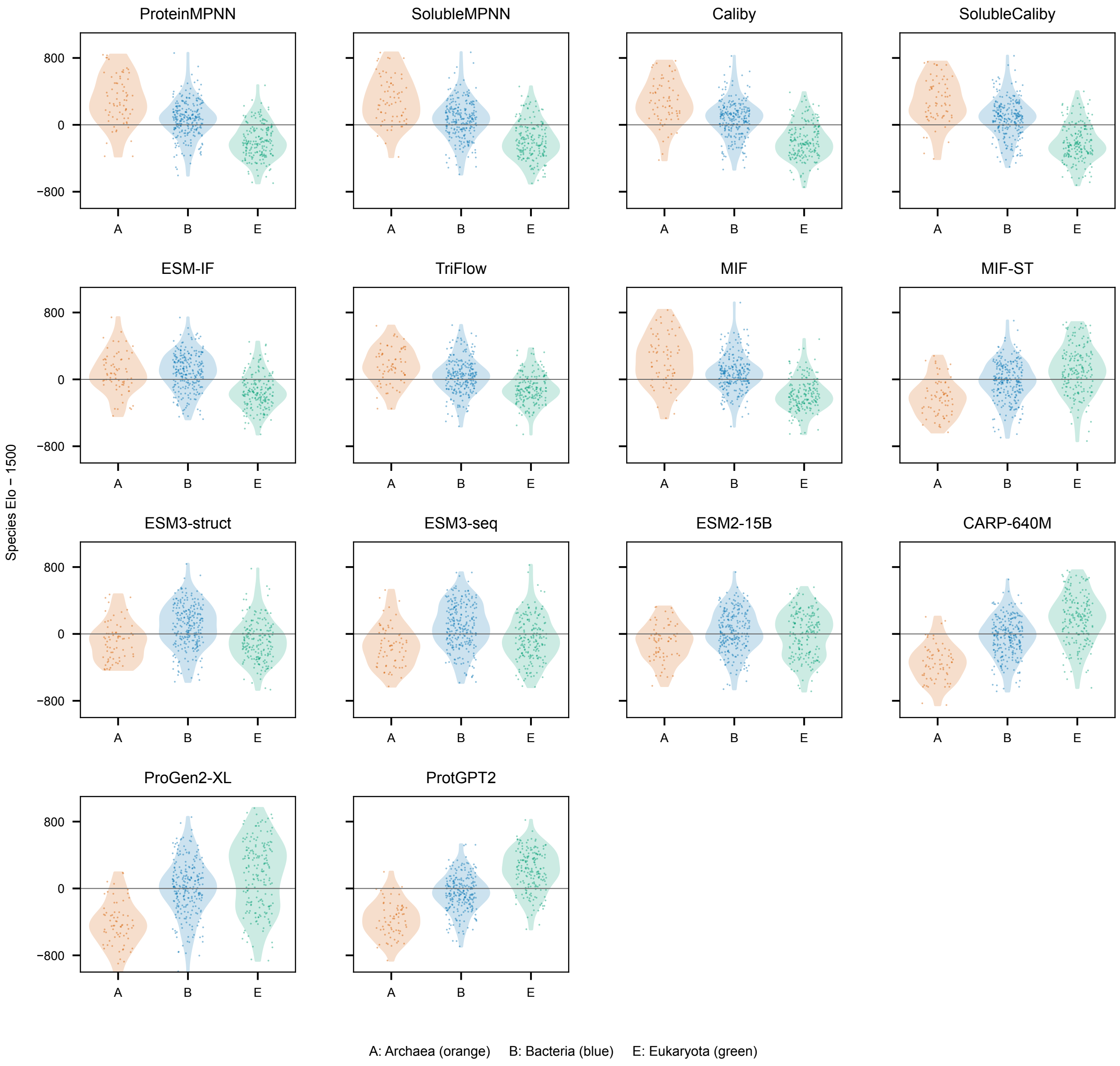

**Figure S4:** Species Elo distributions across all 14 models. Each dot is one species rating relative to the 1,500 baseline, with horizontal jitter for visibility. Violin envelopes show smoothed domain distributions, not confidence intervals. A, Archaea (orange); B, Bacteria (blue); E, Eukaryota (green). Each model has 495 species except ESM2, which has 492 because three species have no available ESM2 scores. Ratings use the unweighted within-family analysis, averaged over 50 match permutations. Source data are provided as a Source Data file.

### S6 Biophysical PCA and model-preference landscapes

#### S6.1 Fourteen-feature PCA

The 14 standardised features are decomposed by principal-component analysis on the natural cohort. PC1 and PC2 explain 23.4% and 16.6% of feature variance, respectively (40.0% cumulative). PC1 separates longer, more hydrophobic, more extended proteins from basic, high-contact-order proteins. PC2 separates heavier, acidic, less stable, more extended proteins from hydrophobic, ordered, compact proteins. Because the plane captures 40% of measured feature variance, it is a coarse biophysical fingerprint rather than a complete representation.

**Table S14:** Feature loadings on the 14-feature PCA. PC1 positive corresponds broadly to larger, hydrophobic, extended proteins; PC1 negative to basic, high-RCO proteins. PC2 positive corresponds to heavier, acidic, unstable, extended proteins; PC2 negative to hydrophobic, ordered, compact proteins.

| Feature | PC1 | PC2 |
|---|---|---|
| sequence length | +0.365 | +0.219 |
| MW per residue | −0.077 | +0.307 |
| isoelectric point | −0.408 | −0.012 |
| acidic-residue fraction | +0.082 | +0.314 |
| basic-residue fraction | −0.458 | +0.264 |
| GRAVY | +0.302 | −0.489 |
| aromaticity | +0.278 | −0.114 |
| instability index | −0.001 | +0.412 |
| proline fraction | +0.033 | +0.087 |
| ordered percentage | −0.005 | −0.209 |
| helix–sheet contrast | +0.258 | −0.040 |
| relative contact order | −0.378 | −0.277 |
| mean C$\beta$ distance | +0.297 | +0.367 |
| surface exposure | −0.102 | −0.061 |

#### S6.2 Taxonomic and functional overlap in the PCA space

The three domains overlap substantially rather than forming separated clusters. The Bhattacharyya overlap of bivariate Gaussian approximations is 95% for Bacteria–Archaea, 91% for Eukaryota–Archaea, and 84% for Eukaryota–Bacteria. Mean PC1 coordinates are −0.66 for Bacteria, −0.17 for Archaea, and +0.66 for Eukaryota. The one-standard-deviation ellipse areas are 6.5, 5.5, and 9.1, respectively. Thus, domain-level trends reflect shifts in density within a shared region rather than taxonomically isolated clusters.

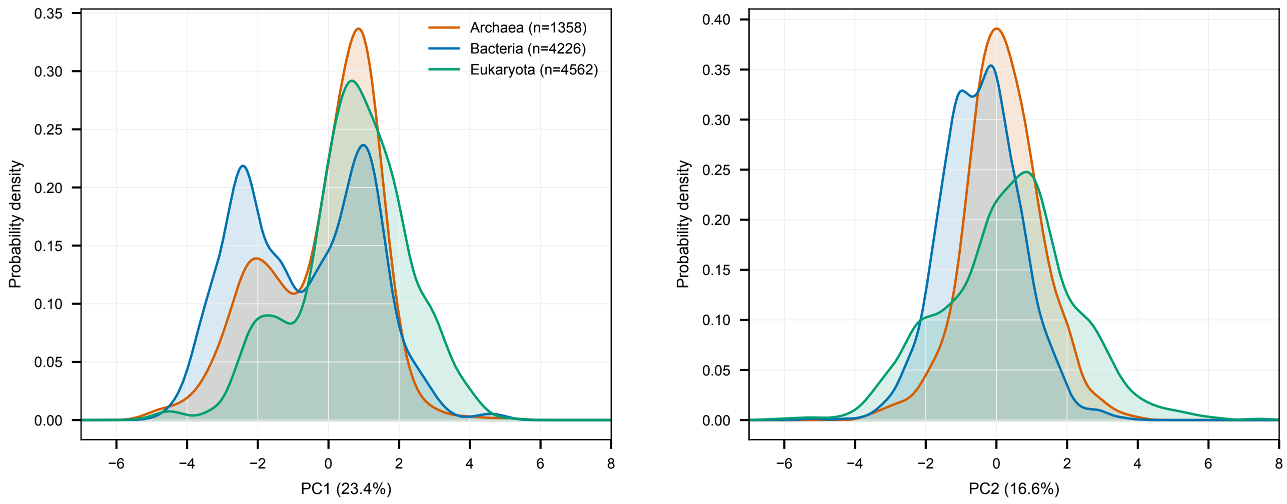

**Figure S5:** PC1 and PC2 distributions by taxonomic domain in the fixed 14-feature natural-protein PCA. Curves are Gaussian kernel-density estimates with Scott's bandwidth, based on 1,358 archaeal, 4,226 bacterial and 4,562 eukaryotic proteins. Orange, blue and green indicate Archaea, Bacteria and Eukaryota. Densities are normalised separately within each domain and show overlap in a shared feature space. Source data include protein coordinates and the displayed density curves. Source data are provided as a Source Data file.

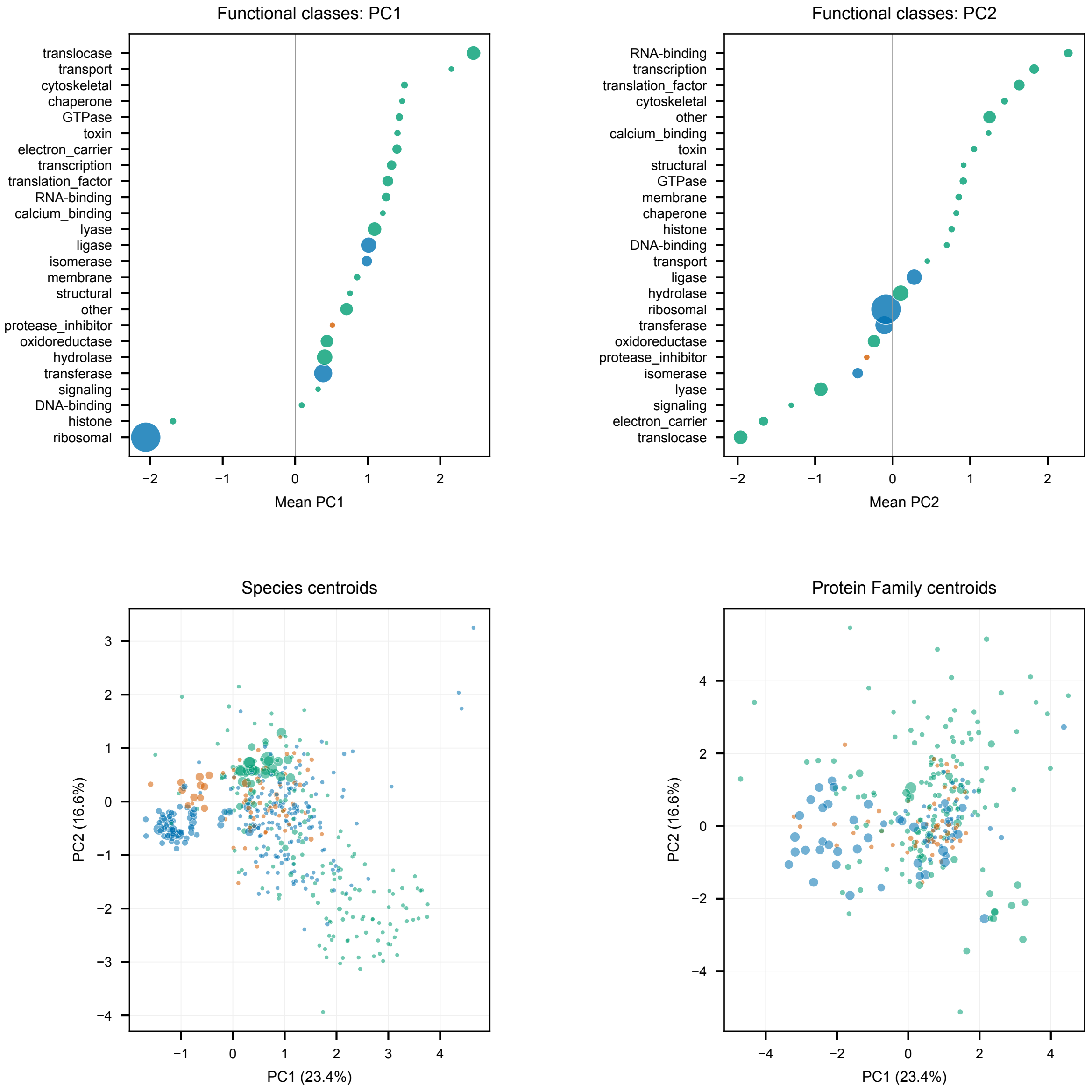

**Figure S6:** Functional and family composition of the biophysical PCA space. Top panels rank broad functional classes separately by their mean PC1 and PC2; the class names shown on each axis follow that panel's ordering. Bottom panels show per-species and protein-family centroids. Each point is one group mean; its area increases with the number of proteins in that group. Colour denotes the most frequent domain in the group: Archaea (orange), Bacteria (blue), or Eukaryota (green). Coordinates, group counts and all underlying protein observations are supplied in Source Data. Source data are provided as a Source Data file.

#### S6.3 Generalised additive model landscapes

For each model, a generalised additive model (GAM) relates z-scored likelihood to PC1 and PC2 using REML in `mgcv`. GAM deviance explained is the fraction of a model's score variation captured by the smooth preference surface; it is distinct from the fraction of feature variance captured by the PCA. Grid cells with fewer than three supporting proteins or outside the observed convex hull are shown faintly for orientation; only supported cells receive full colour and contours and are interpreted. Across models, deviance explained ranges from approximately 9% to 52% and is highest for backbone-conditioned models.

**Table S15:** GAM deviance explained by the 14-feature PC1–PC2 plane. ProteinMPNN uses the `v_48_020` checkpoint. Fine-tuned models are reported for comparison but are not part of the 14-model principal panel. ProGen2-base is an additional checkpoint sensitivity; XL is the representative used in the principal panel.

| Model | GAM deviance explained (%) |
|---|---:|
| ESM-IF | 52.3 |
| SolubleCaliby | 50.9 |
| SolubleMPNN | 50.2 |
| Caliby | 47.5 |
| AcidSecMPNN | 45.1 |
| MIF | 44.3 |
| ProteinMPNN | 43.7 |
| AlkSecMPNN | 43.7 |
| TriFlow | 42.5 |
| ESM3-structure | 40.5 |
| ESM3-sequence | 40.3 |
| MIF-ST | 22.1 |
| CARP-640M | 18.1 |
| ProGen2-base | 14.9 |
| ProtGPT2 | 12.0 |
| ProGen2-XL | 8.7 |
| ESM2-15B | 8.7 |

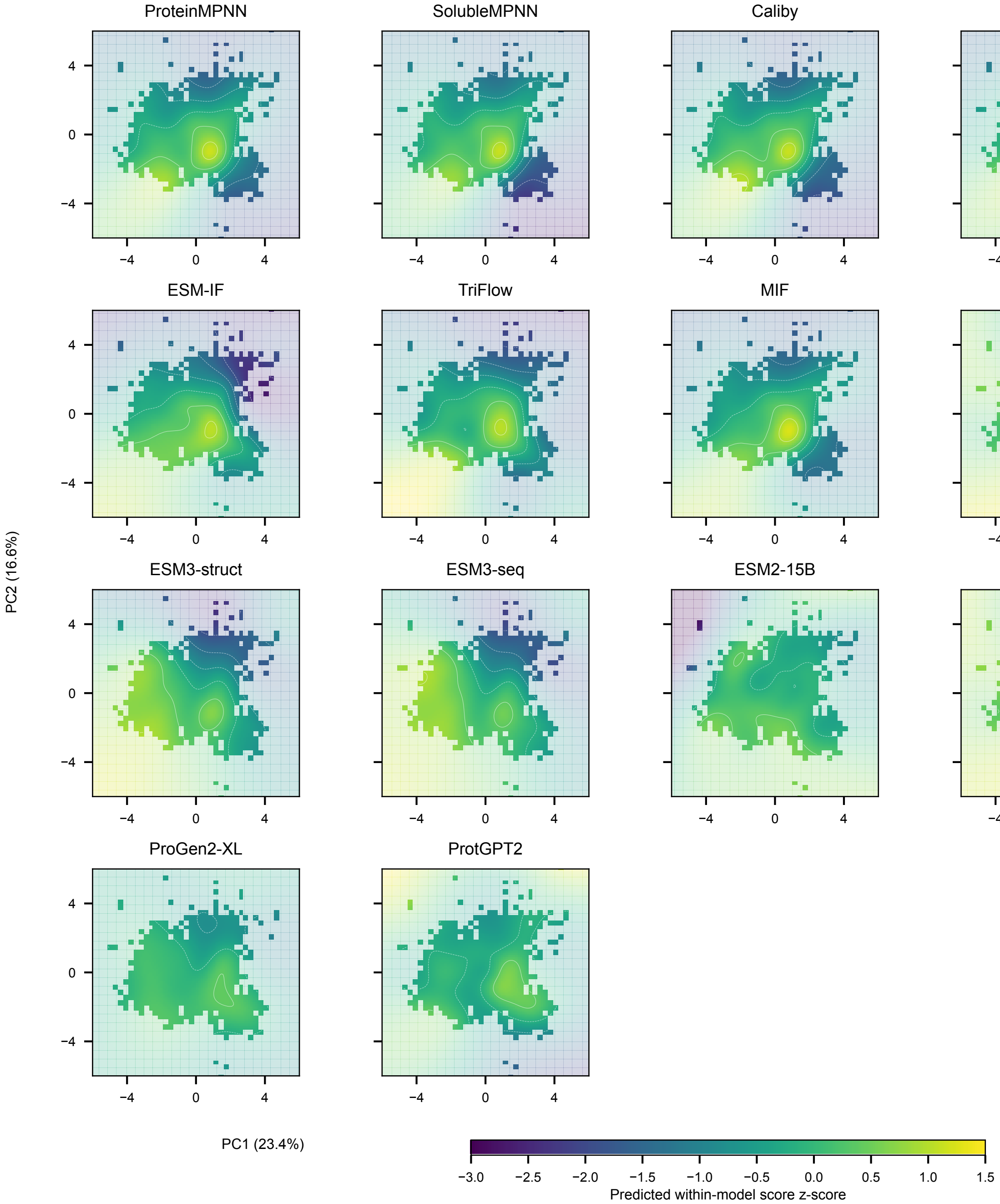

**Figure S7:** Generalised additive model (GAM) score landscapes for the 14 primary models in the global 14-feature PCA frame. Colours show fitted within-model z-scored scores on a shared scale; contours are restricted to supported cells. Predictions with fewer than three supporting proteins or outside the observed convex hull are displayed at 22% opacity as background extrapolation. These faint regions do not establish model preferences in sampled proteins. Fit counts, grids, support flags and display parameters are supplied in Source Data. Source data are provided as a Source Data file.

#### S6.4 Cosine alignment

To test whether a model's preference is organised along an interpretable biophysical direction, we estimate its preference-gradient direction in the PCA plane and calculate its cosine similarity to a reference vector, such as a feature loading or the surface acid–base axis. A large, stable cosine means the model's preference points along that biophysical direction. The same construction is used for the fine-tuning alignment analyses in Sections S9 and S10.

### S7 Property-to-score importance

For each model, we ask which of the 14 measured properties account for why some natural proteins are scored above others. Scores and features are de-meaned within protein family and standardised; species is used as the clustering variable for standard errors. Three measures distribute correlated predictor variance differently:

- Univariate association: $\mathrm{corr}(\tilde{x}_j, \tilde{y})$, which includes variance shared with other features.
- Multivariate coefficient: the joint standardised $\beta_j$, which estimates unique variance but becomes unstable under collinearity. Variance inflation factor is $\mathrm{VIF}_j = 1/(1 - R_j^2)$.
- Johnson relative weight: a decomposition of model $R^2$ that allocates shared variance among correlated features and sums to the fitted $R^2$.

All retained features have VIF below 3. A composition-residualised sensitivity analysis first regresses each score on amino-acid unigram composition and sequence length. Packing and topology remain dominant for backbone-conditioned models, whereas the effects of acidic fraction and GRAVY are reduced.

**Table S16:** Selected ProteinMPNN properties under the within-family importance analysis. Relative weight is expressed as a percentage of model $R^2$; the 14-feature model has $R^2 = 0.29$.

| Property | Univariate | Multivariate $\beta$ | VIF | Relative weight |
|---|---|---|---|---|
| mean C$\beta$ distance | −0.39 | −0.26 | 2.4 | 25.6% |
| relative contact order | +0.33 | +0.13 | 1.9 | 16.6% |
| proline fraction | +0.13 | +0.26 | 1.3 | 12.6% |
| acidic-residue fraction | +0.14 | +0.28 | 2.1 | 10.7% |
| sequence length | −0.25 | +0.01 | 1.6 | 6.8% |
| MW per residue | −0.10 | −0.11 | 2.9 | 2.5% |

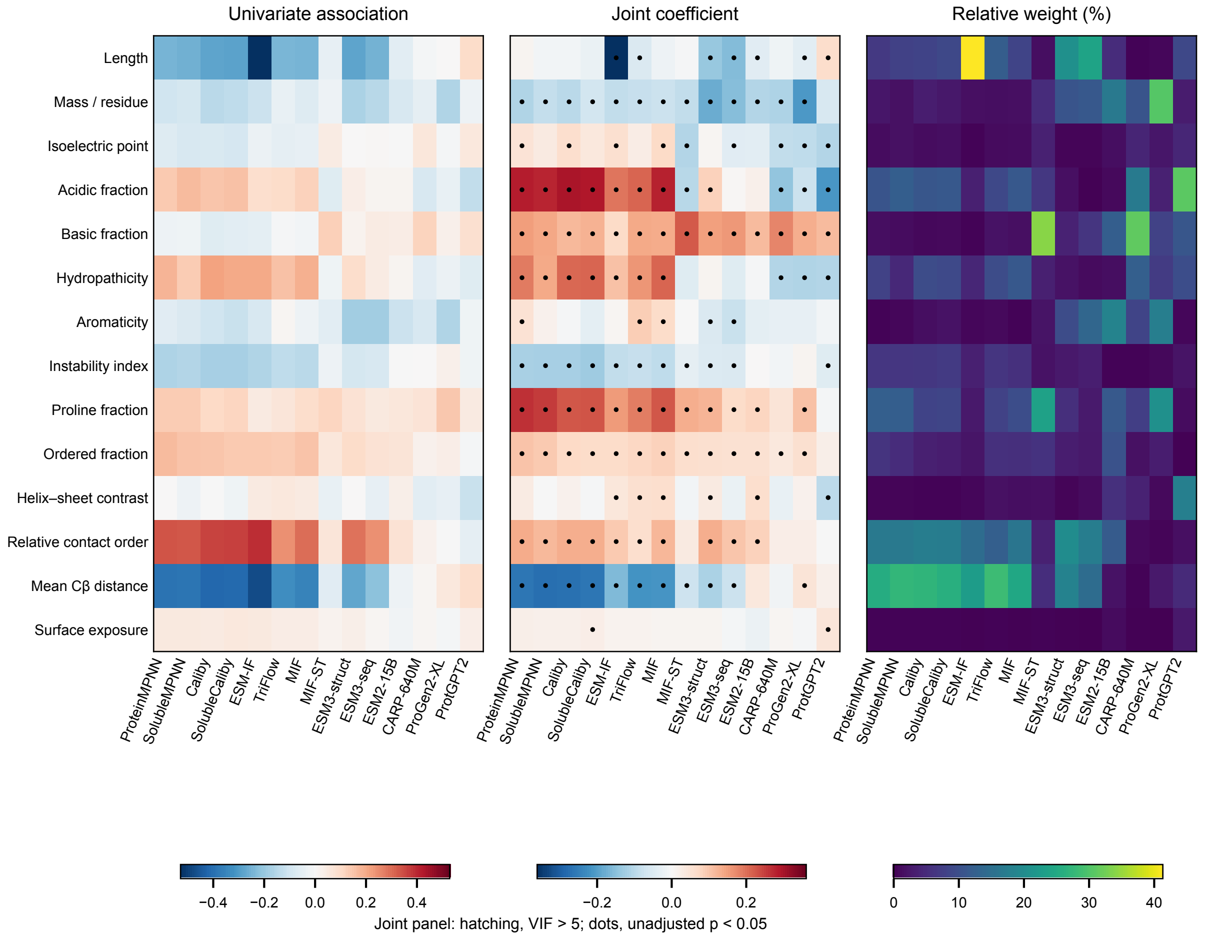

**Figure S8:** Property-to-score importance for the 14 primary models. Heatmaps show within-family univariate standardised coefficients, joint multivariate coefficients, and Johnson relative weights as percentages of within-family $R^2$. Each panel has its own colour scale. Hatching in the joint panel marks a variance inflation factor (VIF) greater than five, and dots mark unadjusted two-sided $p < 0.05$; these are exploratory per-coefficient tests. RCO denotes relative contact order and C$\beta$ denotes the beta carbon. Source data include coefficients, weights, VIFs and p-values. Source data are provided as a Source Data file.

**Table S17:** Johnson relative-weight property importance, expressed as a percentage of each model's within-family $R^2$. ESM3-structure and ESM3-sequence are abbreviated E3s and E3q. PMPNN, ProteinMPNN; SolM, SolubleMPNN; Cal, Caliby; SolC, SolubleCaliby; TriF, TriFlow; ESM2, ESM2-15B; CARP, CARP-640M; PG2, ProGen2-XL; PGP2, ProtGPT2; MW, molecular weight; GRAVY, grand average of hydropathicity. RCO denotes relative contact order.

| Feature | PMPNN | SolM | Cal | SolC | ESM-IF | MIF | MIF-ST | TriF | E3s | ESM2 | E3q | CARP | PG2 | PGP2 |
|---|---|---|---|---|---|---|---|---|---|---|---|---|---|---|
| sequence length | 6.8 | 8.0 | 8.4 | 8.9 | 41.3 | 8.5 | 1.5 | 12.3 | 20.8 | 5.2 | 24.0 | 0.4 | 0.7 | 8.9 |
| MW per residue | 2.5 | 1.9 | 3.2 | 2.6 | 1.7 | 1.8 | 5.3 | 1.6 | 10.8 | 16.6 | 11.3 | 10.8 | 30.4 | 3.2 |
| isoelectric point | 1.4 | 1.8 | 1.4 | 1.6 | 0.5 | 1.4 | 3.8 | 1.2 | 0.5 | 1.5 | 0.6 | 4.3 | 2.5 | 4.4 |
| acidic-residue fraction | 10.7 | 12.3 | 11.1 | 11.4 | 3.7 | 11.4 | 6.7 | 8.9 | 1.6 | 1.0 | 0.3 | 16.9 | 3.7 | 30.9 |
| basic-residue fraction | 1.6 | 1.4 | 0.9 | 1.0 | 0.4 | 1.8 | 34.0 | 2.1 | 3.9 | 12.4 | 6.3 | 31.3 | 8.1 | 11.0 |
| GRAVY | 8.1 | 4.8 | 9.7 | 9.1 | 4.4 | 11.4 | 3.6 | 9.7 | 2.0 | 1.6 | 1.3 | 12.4 | 7.2 | 9.8 |
| aromaticity | 0.5 | 0.8 | 1.5 | 2.1 | 0.8 | 0.3 | 2.2 | 1.1 | 9.6 | 18.5 | 13.6 | 8.5 | 17.7 | 0.7 |
| instability index | 6.5 | 6.5 | 6.7 | 7.1 | 3.7 | 5.7 | 2.1 | 6.3 | 2.9 | 0.3 | 3.4 | 0.4 | 0.9 | 2.2 |
| proline fraction | 12.6 | 12.3 | 8.2 | 8.6 | 2.8 | 9.7 | 23.6 | 8.2 | 5.6 | 11.6 | 2.9 | 7.7 | 20.7 | 1.4 |
| ordered percentage | 6.1 | 5.2 | 3.7 | 3.3 | 2.4 | 5.5 | 6.0 | 5.7 | 3.2 | 10.4 | 2.9 | 1.9 | 3.4 | 0.2 |
| helix–sheet contrast | 0.5 | 0.5 | 0.4 | 0.4 | 0.7 | 1.5 | 1.8 | 1.9 | 0.5 | 5.8 | 1.2 | 3.9 | 1.1 | 17.9 |
| relative contact order | 16.6 | 16.6 | 17.5 | 17.4 | 14.5 | 16.0 | 3.7 | 12.5 | 20.0 | 11.9 | 17.6 | 1.0 | 0.6 | 1.7 |
| mean C$\beta$ distance | 25.6 | 27.4 | 27.0 | 26.1 | 22.9 | 24.7 | 5.3 | 28.3 | 18.4 | 2.0 | 14.7 | 0.4 | 2.8 | 5.0 |
| surface exposure | 0.4 | 0.4 | 0.4 | 0.5 | 0.3 | 0.2 | 0.4 | 0.3 | 0.2 | 1.2 | 0.1 | 0.2 | 0.1 | 2.8 |

##### S7.1 Fine-tuning reshapes local property importance

Re-running the same importance analysis for the two fine-tuned ProteinMPNN models shows that continued training redirects composition-related importance while preserving the structural backbone. AlkSecMPNN increases the acidic-residue weight from 10.7% to 19.5%; AcidSecMPNN reduces it to 3.0% and increases GRAVY from 8.2% to 12.3%. Both retain mean C$\beta$ distance and RCO as the largest terms.

**Table S18:** Relative-weight property importance for base ProteinMPNN and the two fine-tuned models.

| Feature | ProteinMPNN | AlkSecMPNN | AcidSecMPNN |
|---|---|---|---|
| sequence length | 6.8 | 6.5 | 7.5 |
| MW per residue | 2.5 | 5.7 | 6.9 |
| isoelectric point | 1.4 | 2.6 | 0.5 |
| acidic-residue fraction | 10.7 | 19.6 | 3.0 |
| basic-residue fraction | 1.6 | 0.7 | 1.5 |
| GRAVY | 8.1 | 5.3 | 12.2 |
| aromaticity | 0.5 | 2.6 | 1.6 |
| instability index | 6.5 | 6.6 | 6.1 |
| proline fraction | 12.6 | 2.9 | 11.6 |
| ordered percentage | 6.1 | 5.8 | 5.2 |
| helix–sheet contrast | 0.5 | 0.4 | 0.7 |
| relative contact order | 16.6 | 16.8 | 15.6 |
| mean C$\beta$ distance | 25.6 | 23.7 | 26.9 |
| surface exposure | 0.4 | 0.6 | 0.7 |

#### S8 Design-generation shifts and validation controls

##### S8.1 Design templates

The 25 templates span all three domains, lengths 49–633 amino acids, multiple structural classes, and ProteinMPNN Elo preference classes. The cohort contains eight archaeal, seven bacterial, and ten eukaryotic templates.

**Table S19:** Wild-type templates used for chain-A monomer redesign. Rank is the ProteinMPNN Elo preference class.

| UniProt | Domain | Rank | Length | Function | Protein family |
|---|---|---|---|---|---|
| O67786 | Bacteria | Top | 223 | hydrolase | HAD-like hydrolase superfamily |
| Q9Z9J6 | Bacteria | Top | 62 | ribosomal | uL30 |
| Q980U8 | Archaea | Top | 349 | hydrolase | XPG/RAD2 endonuclease |
| O58832 | Archaea | Top | 342 | transferase | DPH1/DPH2 |
| O28542 | Archaea | Top | 340 | oxidoreductase | GAPDH |
| Q8TY15 | Archaea | Top | 361 | ligase | CarA |
| B1X495 | Eukaryota | Top | 103 | translocase | Complex I subunit 4L |
| P0A7R6 | Bacteria | Neutral | 103 | ribosomal | uS10 |
| P60724 | Bacteria | Neutral | 201 | ribosomal | uL4 |
| Q8DLT5 | Bacteria | Neutral | 449 | transferase | GlcNAc-1-P uridyltransferase |
| Q07938 | Eukaryota | Neutral | 337 | transferase | PNP/MTAP phosphorylase |
| Q9N2D5 | Eukaryota | Neutral | 333 | oxidoreductase | GAPDH |
| A9A498 | Archaea | Neutral | 287 | lyase | NnrD/CARKD |
| B1YC30 | Archaea | Neutral | 262 | transferase | RNA polymerase Rpo3/RPB3 |
| A9CIG1 | Bacteria | Low | 101 | ribosomal | uS14 |
| P80505 | Bacteria | Low | 337 | oxidoreductase | GAPDH |
| P02819 | Eukaryota | Low | 49 | structural | Osteocalcin/matrix Gla protein |
| Q4R312 | Eukaryota | Low | 82 | other | DPH3 |
| Q54FJ9 | Eukaryota | Low | 306 | lyase | NnrD/CARKD |
| Q54JL0 | Eukaryota | Low | 633 | oxidoreductase | NADPH diflavin oxidoreductase |
| Q69NP0 | Eukaryota | Low | 93 | other | ATG12 |
| A0A1D8PCG7 | Eukaryota | Low | 87 | ribosomal | eS21 |
| Q6DIY2 | Eukaryota | Low | 179 | oxidoreductase | Acireductone dioxygenase |
| Q6L047 | Archaea | Low | 359 | oxidoreductase | Zn-containing alcohol dehydrogenase |
| G0HQ35 | Archaea | Low | 358 | lyase | Enoyl-CoA hydratase/isomerase |

#### S8.2 Per-property WT-to-design shifts

For each property, the eight designs per template are averaged and paired against that template's WT value, $\Delta_i = \bar{P}_i^{\text{design}} - P_i^{\text{WT}}$. The paired standardised effect is

$$d_z = \frac{\text{mean}(\Delta)}{\text{sd}(\Delta)}. \quad \text{(S11)}$$

Two-sided Wilcoxon signed-rank tests use a normal approximation with continuity correction and exclude zero differences. Paired differences are rounded to 12 decimal places for ranking only, to avoid spurious ranks from floating-point representations of equal composition shifts; means and $d_z$ use unrounded values. Benjamini–Hochberg correction is applied jointly across all 84 model–property tests (seven models and 12 varying properties). Chain length is fixed by construction. The global surface-exposure definition is backbone-derived and is not treated as a freely varying sequence property in the primary 12-property shift summary.

Designs are folded with single-sequence ColabFold (five models, three recycles) to obtain structural features. The natural-protein PCA uses AlphaFold-DB models, so per-feature mean offsets measured on the 25 WTs folded by both pipelines place designs in the AlphaFold-DB coordinate system. This offset does not affect within-predictor WT-to-design differences.

Five properties have significant shifts in at least four of the six non-MIF-ST design models: pI, acidic-residue fraction, aromaticity, proline fraction, and helix–sheet contrast. Relative contact order shifts significantly in three of these models. The models tend to generate more acidic compositions, more proline, greater helix–sheet contrast, lower aromaticity, and, for ProteinMPNN- and Caliby-family models, greater contact order. MIF-ST differs from the other models and is considered separately in Section S11.

**Table S20:** Wild-type-to-design shifts for the 12 properties that vary in the primary fixed-backbone analysis. Values are paired effect sizes $d_z$ across 25 templates, each represented by the mean of eight designs. Asterisks denote Benjamini–Hochberg $q < 0.05$ across all 84 model–property tests, using two-sided normal-approximation Wilcoxon signed-rank tests with continuity correction.

| Property | ProteinMPNN | SolubleMPNN | Caliby | SolubleCaliby | ESM-IF | MIF | MIF-ST |
|---|---|---|---|---|---|---|---|
| Proline fraction | $+1.37^*$ | $+1.01^*$ | $+2.02^*$ | $+1.75^*$ | $-0.09$ | $+0.47$ | $-0.28$ |
| Acidic fraction | $+0.38$ | $+0.89^*$ | $+0.80^*$ | $+0.98^*$ | $+0.83^*$ | $+0.61^*$ | $-0.70^*$ |
| Helix–sheet contrast | $+0.75^*$ | $+1.01^*$ | $+0.65^*$ | $+0.64^*$ | $+0.52^*$ | $+0.81^*$ | $+0.78^*$ |
| Isoelectric point | $-0.47$ | $-0.86^*$ | $-0.59^*$ | $-0.83^*$ | $-0.44$ | $-0.71^*$ | $+0.96^*$ |
| Aromaticity | $-0.70^*$ | $-0.80^*$ | $-0.76^*$ | $-0.76^*$ | $-0.42$ | $-0.61^*$ | $+0.54^*$ |
| Relative contact order | $+0.32$ | $+0.49^*$ | $+0.61^*$ | $+0.58^*$ | $-0.12$ | $-0.15$ | $-0.52^*$ |
| Mean C$\beta$ distance | $-0.20$ | $-0.20$ | $-0.46$ | $-0.45$ | $-0.01$ | $-0.15$ | $+0.77^*$ |
| Basic fraction | $+0.00$ | $+0.18$ | $+0.27$ | $+0.30$ | $+0.44$ | $+0.09$ | $+0.64^*$ |
| MW per residue | $-0.60^*$ | $-0.15$ | $-0.12$ | $-0.04$ | $-0.18$ | $-0.58^*$ | $-0.15$ |
| Instability index | $+0.06$ | $+0.21$ | $-0.15$ | $-0.05$ | $+0.11$ | $-0.15$ | $-1.02^*$ |
| GRAVY | $+0.26$ | $-0.23$ | $-0.17$ | $-0.24$ | $-0.11$ | $+0.12$ | $-0.19$ |
| Ordered percentage | $+0.11$ | $+0.07$ | $+0.21$ | $+0.21$ | $+0.21$ | $+0.02$ | $-0.24$ |

#### S8.3 Functional-residue recovery

For each template, the union of UniProt active-site, binding-site, metal-binding, site, DNA-binding, and nucleotide-binding positions defines the functional set. Fifteen templates contain at least one curated functional position and have a canonical length matching the WT design input. Designs contain no insertions or deletions, so annotations map directly to design positions. For each model, functional-site recovery is compared with recovery at all other positions using a two-sided Wilcoxon signed-rank test across templates.

No model preferentially preserves curated functional residues. MIF-ST is the only significant result and recovers functional positions below background. Differential retention of annotated catalytic and binding residues therefore does not explain the shared shifts among the other models. The analysis does not cover all protein interfaces because consistent interface annotation is sparse across this broad cohort.

**Table S21:** Recovery at curated functional positions and background positions. Replicates are averaged within each of 15 templates before the paired Wilcoxon test.

| Model | $r_{\text{func}}$ | $r_{\text{bg}}$ | Difference | $p$ |
|---|---|---|---|---|
| ESM-IF | 0.71 | 0.64 | $+0.075$ | 0.30 |
| MIF | 0.46 | 0.50 | $-0.036$ | 0.25 |
| ProteinMPNN | 0.44 | 0.49 | $-0.049$ | 0.39 |
| AlkSecMPNN | 0.45 | 0.55 | $-0.095$ | 0.12 |
| AcidSecMPNN | 0.42 | 0.53 | $-0.112$ | 0.14 |
| SolubleMPNN | 0.44 | 0.50 | $-0.055$ | 0.39 |
| SolubleCaliby | 0.36 | 0.44 | $-0.078$ | 0.23 |
| Caliby | 0.36 | 0.44 | $-0.079$ | 0.25 |
| MIF-ST | 0.20 | 0.35 | $-0.158$ | $1.8 \times 10^{-4}$ |

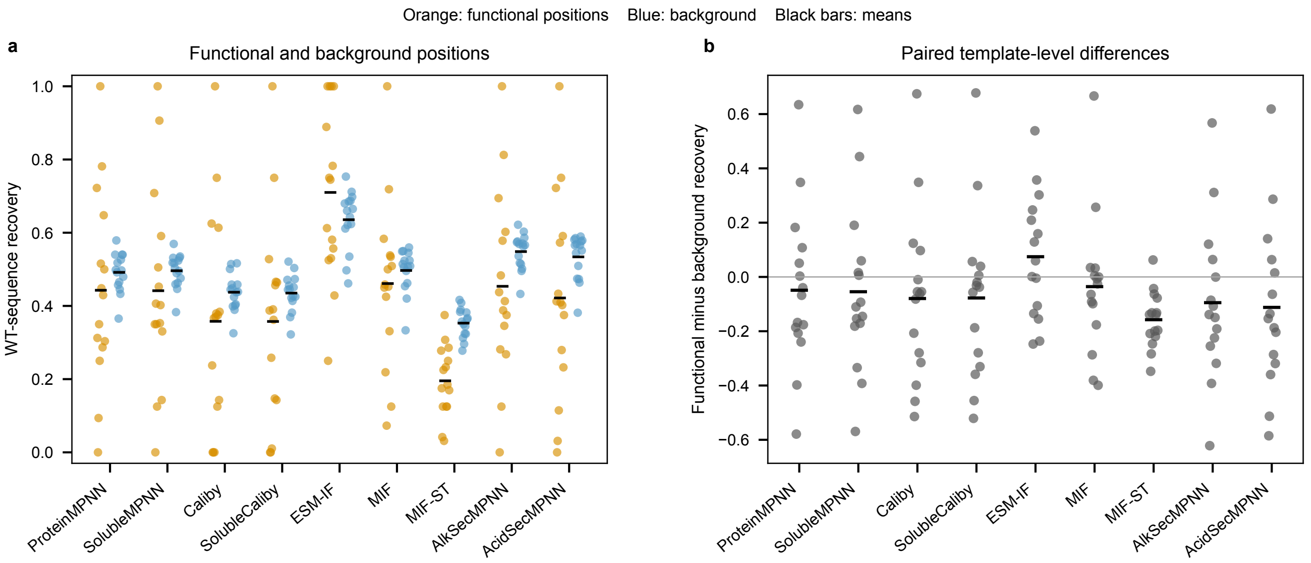

**Figure S9:** Functional-residue recovery across 15 annotated templates per model. a, Each point is the mean recovery across eight designs for one template, at curated UniProt functional positions (orange) or background positions (blue); black bars show means across templates. b, Paired functional-minus-background differences for the same 15 templates, with black bars showing means. All nine plotted models are included in Supplementary Table S21; the seven primary models are also summarised in Supplementary Table S25. Two-sided paired Wilcoxon tests use templates as the independent units; the reported p-values are unadjusted. Only MIF-ST has a significant difference at $p<0.05$, recovering functional positions below background. Source data are provided as a Source Data file.

##### S8.4 Score–generation coherence and importance non-transfer

Across the 12 varying properties, the signed scoring reward correlates positively with the signed design push for all six non-MIF-ST design models (mean Pearson $r \approx 0.41$). ProteinMPNN shows the strongest association ($r = 0.50$; Spearman $\rho = 0.64$). Thus, scoring preferences and design shifts reflect a shared tendency, although the natural-cohort property-importance ranking does not transfer cleanly to a model's own high-likelihood designs. The design populations occupy a narrow region in which within-design property gradients are weak, even when biophysical features explain substantial variance in self-score.

#### S9 ProteinMPNN fine-tuning on extremophile secretomes

##### S9.1 Rationale and cohorts

Continued training tests whether a biologically structured change in the training distribution can redirect ProteinMPNN's design preference without overwriting backbone compatibility. Cases are secreted proteins from organisms with defined environmental growth pH, not proteins selected directly by charge or pI. Alkaliphile organisms span approximately pH 9–11 and acidophile organisms pH 1–4. Secreted proteins were selected because extracellular pH acts directly on their surfaces. Each case was matched one-to-one to a neutralophile-secretome protein using Pfam where possible, then enzyme class and length. Sequences were clustered at 40% identity and split 70/15/15 so no cross-split pair exceeded 40% identity. The alkaliphile cohort contained approximately 250 matched training pairs and the acidophile cohort approximately 75.

##### S9.2 Continued-training protocol

Each arm produced an extremophile model and a matched neutralophile control trained under the same protocol. Primary runs initialise from ProteinMPNN `v_48_020`; checkpoint sensitivity runs initialise from `v_48_002`. Training continues the backbone-conditioned negative-log-likelihood objective with 0.2 Åbackbone noise. The selected checkpoint is the latest whose validation native- sequence recovery remains within three percentage points of the base model.

##### S9.3 Held-out surface-chemistry shifts

Fine-tuned models and their bases redesigned held-out neutralophile backbones using eight designs per backbone at $T = 0.1$. Surface residues have relative solvent accessibility at least 0.25 under Shrake–Rupley SASA with Tien normalisation. A surface acid–base PCA provides a low-dimensional summary. It is fitted after sample-standardisation of four features (surface acidic, basic and ionisable residue fractions, and surface net charge) across 1,225 records: 25 WTs and 200 designs from each of six primary or sensitivity models. PC1 is oriented so positive means higher surface acidic-residue fraction. The primary and sensitivity comparisons are evaluated separately after projection into this fixed frame.

Relative to base ProteinMPNN, AlkSecMPNN shifts designs toward more acidic surfaces ($\Delta$PC1 $= +1.94$), while AcidSecMPNN shifts in the opposite direction ($\Delta$PC1 $= -1.00$). Direct features show that AlkSecMPNN raises surface acidic fraction and lowers net charge, charge per residue, and pI. AcidSecMPNN raises net charge and pI while reducing surface acidic fraction, indicating that its shift occurs mainly through loss of acidic residues rather than addition of basic residues.

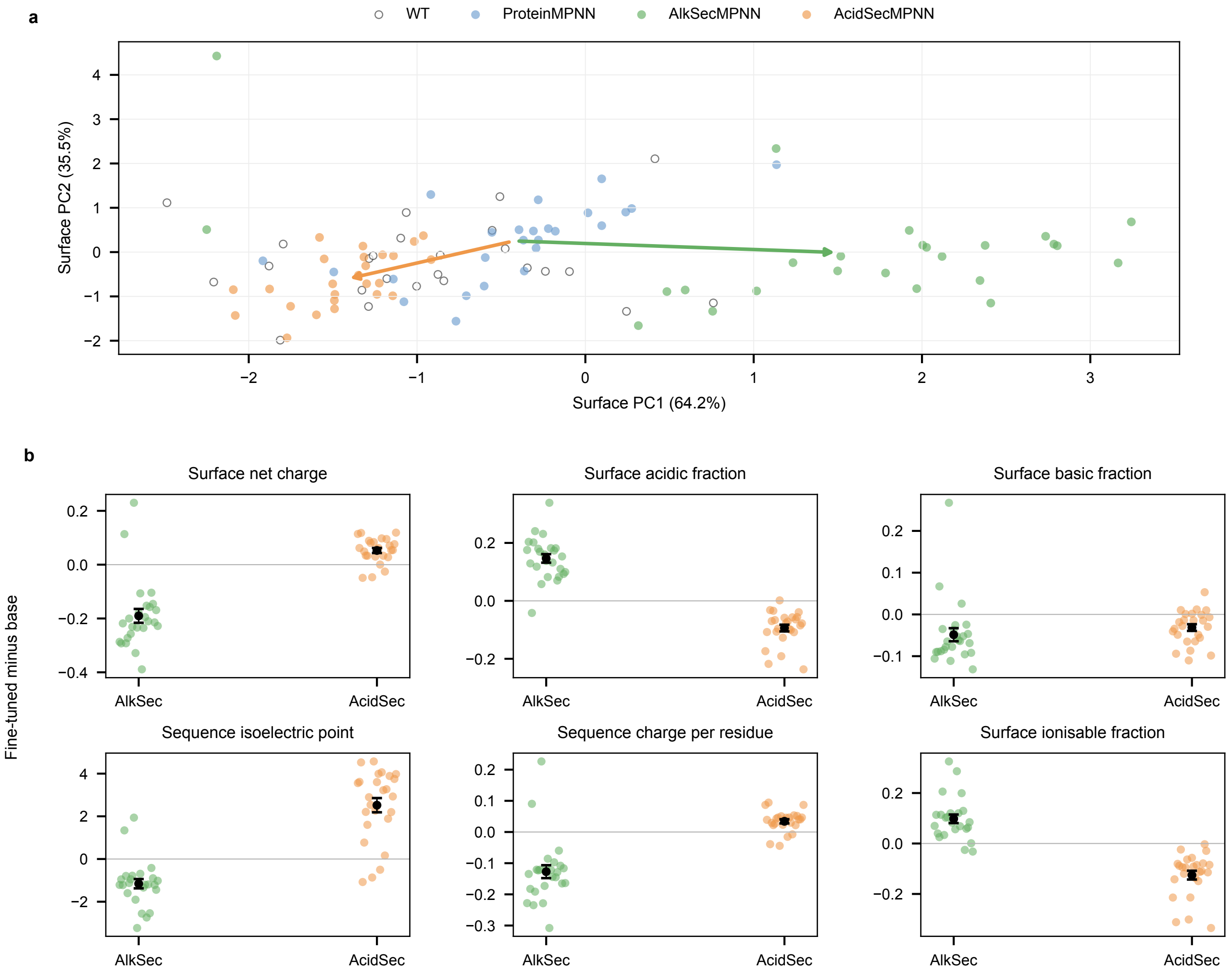

**Figure S10:** Fine-tuning redirects design surface chemistry across 25 templates. a, Each point is a WT or a model-specific mean of eight designs for one template in the four-feature surface acid–base PCA; arrows connect the base-model centroid to each fine-tuned centroid. The fit uses 1,225 WT/design records across primary and sensitivity checkpoints, as described in Methods; the plotted models are the primary v_48_020 runs. b, Each coloured point is a template-level fine-tuned-minus-base difference for a direct surface or sequence feature (n=25 per model); black points and error bars show the mean and standard error of the mean. Positive values indicate an increase relative to base. The corresponding paired tests and Holm-adjusted p-values are supplied in the Source Data. Source data are provided as a Source Data file.

**Table S22:** Base-relative direct surface-feature shifts for the primary `v_48_020` models. Surface residues have relative solvent-accessible surface area (SASA) at least 0.25.

| Feature | AlkSecMPNN | AcidSecMPNN |
|---|---:|---:|
| Surface acidic fraction | +0.147 | −0.094 |
| Surface basic fraction | −0.049 | −0.031 |
| Surface ionisable fraction | +0.098 | −0.125 |
| Surface net charge | −0.190 | +0.053 |
| Charge per residue | −0.127 | +0.034 |
| Isoelectric point | −1.16 | +2.52 |

#### S9.4 Matched controls and design quality

Matched neutralophile controls indicate that the alkaliphile arm is the cleaner perturbation: its control shifts in the opposite surface-charge direction. The acidophile arm is directionally consistent but more confounded because its neutralophile control partly follows the same direction. Native-sequence recovery changes by −1.4 and +0.3 percentage points for the two arms, while entropy, duplicate rate, and pairwise identity remain comparable with the base model.

#### S9.5 Single-sequence structural self-consistency

Designs and WT sequences were refolded with single-sequence ColabFold (five models, three recycles) and aligned to the input AlphaFold-DB backbone. Structural agreement is measured by self-consistency TM-score, C$\alpha$ RMSD, and pLDDT, with paired fine-tuned-versus-base tests across templates. AlkSecMPNN is indistinguishable from base (TM-score 0.51 versus 0.54; difference −0.02, not significant). AcidSecMPNN shows a small reduction (TM-score 0.45; difference −0.09, $p < 0.001$; pLDDT lower by approximately six), but both fine-tuned arms exceed the WT single-sequence control (TM-score 0.38). These checks support structural compatibility but do not replace experimental validation.

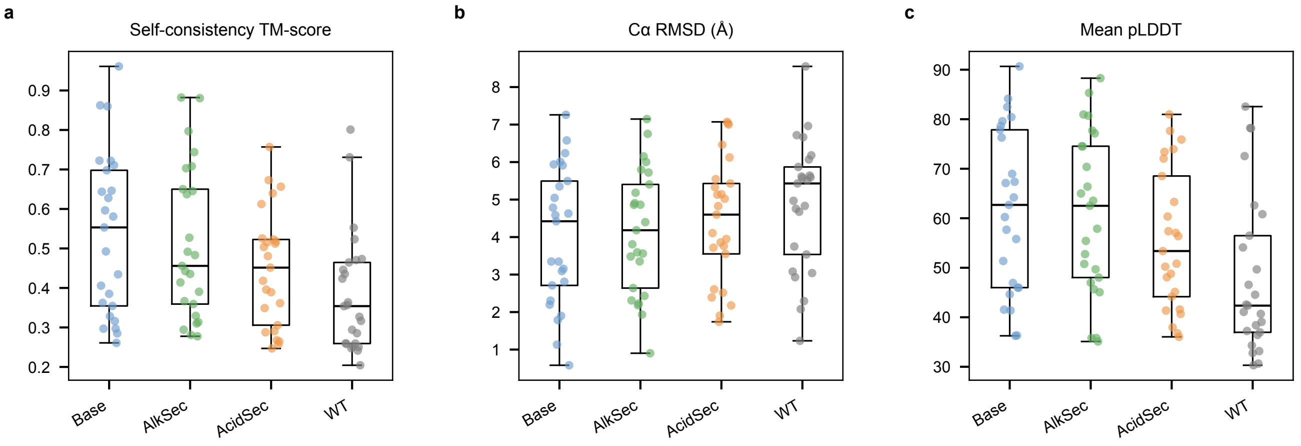

**Figure S11:** Structural self-consistency across 25 templates. Panels show a, TM-score normalised by the AlphaFold-DB input reference, calculated with TM-align through tmtools; b, C$\alpha$ root mean square deviation (RMSD) in angstroms; and c, mean predicted local distance difference test (pLDDT) confidence. Each point is a template-level mean across eight single-sequence ColabFold designs; WT controls have one refold per template (n=25 per group). Boxes span the first to third quartiles, central lines mark medians, and whiskers extend to the most extreme observations within 1.5 times the interquartile range; all points are shown. Fine-tuned-versus-base tests pair template means, as reported in the Source Data. Source data are provided as a Source Data file.

#### S9.6 Global taxonomic structure is preserved

Fine-tuning does not overwrite the broader taxonomic preference: the Archaea–Eukaryota Elo gap is +496 for base ProteinMPNN, +553 for AlkSecMPNN, and +417 for AcidSecMPNN. Continued training therefore redirects a local surface-chemistry tendency while retaining the global structure-conditioned taxonomic pattern.

### S10 Sequence-model malleability control

#### S10.1 ESM2-35M continued training

To test whether the design shift is specific to a structure-conditioned model, ESM2-35M (`esm2_t12_35M_UR50D`) was fine-tuned by continued masked-language-model training on the same alkaliphile-secretome cases and cluster-disjoint splits used for AlkSecMPNN. A matched neutralophile-control fine-tune separates cohort-specific effects from generic continued training on secreted proteins. Training uses 15% masking, AdamW, and 30 epochs; a schedule sweep tests sensitivity to training amount.

#### S10.2 Matched surface-only redesign

Both ESM2-35M and ProteinMPNN redesign the same exposed positions on ten held-out neutralophile secreted targets, with buried residues fixed at WT. ProteinMPNN uses fixed-position generation; ESM2-35M uses Gibbs-style iterative masked infilling. Both generate eight sequences per target at $T = 0.1$. The response is surface net charge, $(K + R - D - E)/n_{\text{surface}}$, and the fine-tuning steer is the extremophile model minus its matched neutralophile control.

Fine-tuning steers both models toward more acidic surfaces, but ProteinMPNN moves approximately three times as far. AlkSecMPNN changes surface net charge by $-0.232 \pm 0.037$ and shifts all ten targets in the acidic direction. AlkSecESM2-35M changes it by $-0.075 \pm 0.021$ and shifts nine of ten. This comparison establishes that a sequence-only model is malleable under the same data intervention, while the structure-conditioned model produces a larger change on the matched generation task.

**Table S23:** Matched surface-only redesign of ten secreted targets. Values are mean fine-tuned-minus-control surface-net-charge shifts ± the standard error of the mean (s.e.m.); more negative means more acidic.

| Model | Surface-net-charge shift | Targets shifted acidic |
|---|---|---|
| ProteinMPNN (structure-conditioned) | $-0.232 \pm 0.037$ | 10/10 |
| ESM2-35M (sequence-only) | $-0.075 \pm 0.021$ | 9/10 |

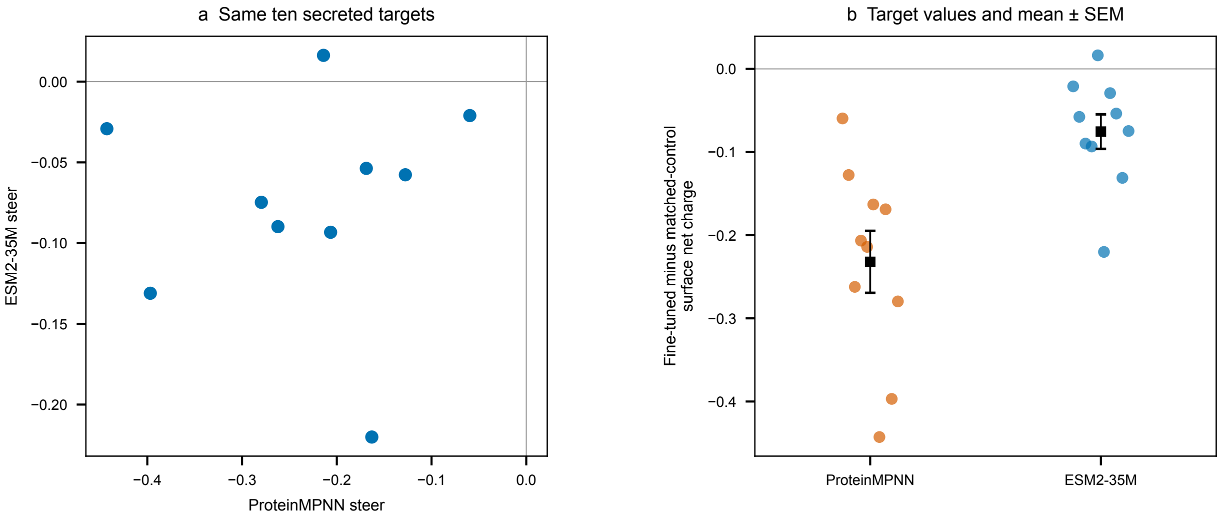

**Figure S12:** Matched sequence- versus structure-conditioned malleability on the same ten secreted targets. Steer is the fine-tuned-minus-matched-neutralophile-control change in surface net charge; more negative values indicate a larger acidic shift. a, Each point pairs the ProteinMPNN and ESM2-35M steer for one target. b, All ten target values are shown per model; black squares and error bars show the mean and standard error of the mean (SEM) across targets (n=10 per model). Source data are provided as a Source Data file.

#### S10.3 Per-position sequence-model preference shift

The per-position probability difference shows how ESM2 fine-tuning raises preference for acidic residues before generation. The effect is concentrated at exposed positions, consistent with the aggregate surface-net-charge shift.

supplementary_figure_13.png

**Figure S13:** Per-position ESM2 preference shift for a representative target. The upper heatmap shows fine-tuned-minus-base amino-acid probability at each position; red means greater preference after fine-tuning. The lower panel shows acidic propensity, $P(\mathrm{D}) + P(\mathrm{E}) - P(\mathrm{K}) - P(\mathrm{R})$, under base, AlkSec, and matched neutralophile-control ESM2-35M models for Q9Z9J6 (62 residues). Shading marks the 42 surface positions with relative solvent-accessible surface area (SASA) at least 0.25. Source data are provided as a Source Data file.

#### S11 Sequence-pretrained boundary cases

MIF-ST is best interpreted as a sequence-pretrained boundary case rather than excluded as a generic outlier. MIF and MIF-ST share a masked inverse-folding architecture and structural training data; MIF-ST additionally incorporates CARP-640M sequence representations. This change moves MIF-ST toward the sequence-like regime on several independent measures:

- residual species variance is 15.9%, attenuation 33%, and within-family retention 0.75, compared with 3.4%, 87%, and 0.31 for MIF;
- its Archaea–Eukaryota Elo gap is negative (−338), whereas MIF's is positive (+412);
- its design-shift profile is negatively correlated with the other structural-input models;
- it is the only model to recover curated functional residues significantly below background.

ESM3-sequence is a boundary case in the opposite direction: it is scored without supplied structure but shows only 5.2% residual species variance and 84% attenuation. Together, these cases show that the three scoring contexts describe dominant tendencies rather than exhaustive architectural categories. Sequence-derived pre-training can make a model with structural input behave sequence-like, while a multimodal architecture scored in sequence-only mode need not reproduce the residual taxonomic preference of classical protein language models.

#### S12 Tables supporting the main article

This section reports additional biophysical, functional-recovery and surface-chemistry summaries, together with dataset and feature definitions. The species-signal and domain Elo summaries are presented in main Tables 1 and 2.

**Table S24:** Measured biophysical features explain more unique score variance in backbone-conditioned models than in classical sequence-only language models. The biophysics partial $R^2$ is $(R^2_{\text{Full}} - R^2_{\text{Family+Species}})/(1 - R^2_{\text{Family+Species}})$: the fraction of variance remaining after species and family adjustment that is explained by the 14 biophysical features. This differs from the total-variance increment $\Delta_{\text{bio}}$ defined above. Scoring contexts are defined in Table 1 of the main article. Values show the range across models within each context. Full model-level values are reported in Supplementary Table S6.

| Scoring context | Number of models | Biophysics partial $R^2$ |
|---|---|---|
| Backbone-conditioned | 6 | 0.11–0.34 |
| Structure + native sequence context | 3 | 0.07–0.15 |
| Sequence-only | 5 | 0.03–0.11 |

**Table S25:** Recovery at annotated functional positions vs background ($n = 15$ templates; two-sided Wilcoxon signed-rank). No model preferentially preserves functional residues; only MIF-ST differs significantly (under-recovering).

| Model | $r_{\text{func}}$ | $r_{\text{bg}}$ | $\Delta$ | $p$ |
|---|---|---|---|---|
| ESM-IF | 0.71 | 0.64 | +0.075 | 0.30 |
| MIF | 0.46 | 0.50 | −0.036 | 0.25 |
| ProteinMPNN | 0.44 | 0.49 | −0.049 | 0.39 |
| SolubleMPNN | 0.44 | 0.50 | −0.055 | 0.39 |
| SolubleCaliby | 0.36 | 0.44 | −0.078 | 0.23 |
| Caliby | 0.36 | 0.44 | −0.079 | 0.25 |
| MIF-ST | 0.20 | 0.35 | −0.158 | $1.8 \times 10^{-4}$ |

**Table S26:** Base-relative design shift along the surface acid–base PCA (PC1 oriented so positive = more acidic, lower-charge surface).

| Comparison | $\Delta$PC1 vs. base | Interpretation |
|---|---|---|
| AlkSecMPNN vs ProteinMPNN | +1.94 | more acidic surface |
| AcidSecMPNN vs ProteinMPNN | −1.00 | less acidic / higher-charge surface |

**Table S27:** Surface-feature shifts and matched neutralophile controls. A, Direct base-relative shifts for the common 25-template design set (`v_48_020`; protein-level means; surface = relative SASA $\geq 0.25$). B, Mean changes in surface net charge on held-out neutralophile backbones from each arm, where $n$ is the number of backbones. Each comparison is calculated as the first named model minus the second on the same backbones.

*A. Direct surface-feature shifts on the common 25-template design set*

| Feature | AlkSecMPNN | AcidSecMPNN |
|---|---|---|
| Surface acidic fraction | +0.147 | −0.094 |
| Surface basic fraction | −0.049 | −0.031 |
| Surface ionizable fraction | +0.098 | −0.125 |
| Surface net charge | −0.190 | +0.053 |
| Charge per residue | −0.127 | +0.034 |
| Isoelectric point | −1.16 | +2.52 |

*B. Matched-control evaluation of surface net charge*

| Training arm | Held-out backbones ($n$) | Extremophile minus base | Neutralophile control minus base | Extremophile minus control |
|---|---|---|---|---|
| Alkaliphile | 53 | −0.1400 | +0.0957 | −0.2358 |
| Acidophile | 14 | +0.1018 | +0.0770 | +0.0248 |

**Table S28:** Main Dataset Composition. This dataset was used for all analyses unless otherwise specified.

| Metric | Archaea | Bacteria | Eukaryota |
|---|---|---|---|
| Species | 68 | 242 | 185 |
| Proteins | 1,358 (13.4%) | 4,226 (41.7%) | 4,562 (45.0%) |
| Protein families | | | |
| Total families | | | 281 |
| Three-domain families | | | 74 |
| Two-domain families | | | 88 |
| Single-domain families | | | 119 |
| Protein characteristics | | | |
| Sequence length range | | | 24–2,531 amino acids |
| Mean pLDDT | | | 89.7 ± 7.9 |
| Mean isoelectric point | | | 7.6 ± 2.3 |
| Mean basic residue fraction | | | 0.156 ± 0.055 |

**Table S29:** The 14 modelling features (9 sequence, 5 structure). Net charge at pH 7 and small-residue fraction are also computed and reported in the design-shift analysis but excluded from the PCA/variance-decomposition/property-importance set as collinear with isoelectric point and MW per residue.

| Sequence (9) | Structure (5) |
|---|---|
| sequence length | ordered percent (helix% + sheet%) |
| MW per residue | helix−sheet contrast (helix% − sheet%) |
| isoelectric point | relative contact order (RCO) |
| acidic-residue fraction | mean C$\beta$ distance |
| basic-residue fraction | surface exposure |
| GRAVY (mean hydropathy) | |
| aromaticity | |
| instability index | |
| proline fraction | |
